# A Comparative Nanomechanical Study of Antibody and Nanobody Binding to SARS-CoV-2 Variants

**DOI:** 10.1101/2025.09.26.672567

**Authors:** Luis F. Cofas-Vargas, Gustavo E. Olivos-Ramirez, Siewert J. Marrink, Adolfo B. Poma

**Affiliations:** Biosystems and Soft Matter Division, Institute of Fundamental Technological Research, Polish Academy of Sciences, ul. Pawińskiego 5B, 02-106 Warsaw, Poland; Groningen Biomolecular Sciences and Biotechnology Institute, University of Groningen, Nijenborgh 7, 9747 AG Groningen, The Netherlands

## Abstract

The receptor-binding domain of the SARS-CoV-2 spike protein is the principal target of neutralizing antibodies (Abs) and nanobodies (Nbs). Although their thermodynamic binding properties have been extensively characterized, their stability under mechanical force remains less understood. Here, we perform a comparative nanomechanical analysis of three Abs (PDI-231, S2X259, and R1-32) and three Nbs (R14, C1, and n3113.1) bound to the RBD from the WT strain and the Omicron BA.4 and JN.1 variants. Using coarse-grained steered molecular dynamics within the GōMartini 3 framework, we identified distinct force–response behaviors shaped by epitope topology, binding architecture, and variant-specific mutations. Ab/RBD dissociation was characterized by asymmetric rupture events, variant-dependent unfolding of RBD segments, and occasional deformation of antibody constant domains. Analysis of single-chain systems revealed that the heavy chain acts as the main load-bearing element, while the light chain sustains a consistent but weaker mechanical response. For the two-chain Ab system, the cooperative action of both chains enhances stability, enabling complexes to withstand rupture forces in the range of 500 pN. By contrast, Nb/RBD complexes dissociated primarily through rigid-body mechanisms, transmitting force more directly to the RBD interface with minimal structural disruption. Together, these results demonstrate that mechanical resilience emerges from immune complex topology and inter-chain cooperation, providing complementary insights beyond affinity into the design of therapeutics resilient to viral evolution.

## 1 Introduction

The coronavirus disease (COVID-19) pandemic, caused by the betacoronavirus severe acute respiratory coronavirus 2 (SARS-CoV-2), has resulted in millions of infections and deaths worldwide ^1^. SARS-CoV-2 encodes four structural proteins, including the trimeric spike (S) glycoprotein, which facilitates virus entry through recognition of the human angiotensin-converting enzyme 2 (hACE2) ^2^. Each S monomer is composed of a N-terminal S1, responsible for receptor binding, and a C-terminal S2 subunit, which drives membrane fusion^3^. Within S1, the receptor-binding domain (RBD; residue range R319-F541), sits at the apex of the trimer and contains the receptor-binding motif (RBM) comprising residues S438-Q506, which directly engages hACE2 ^4,5^.

The RBD is a primary target of human neutralizing antibodies (Abs) and engineered nanobodies (Nbs) because of its crucial role in host recognition ^6^. Mutations within the RBM have been key drivers in the emergence of SARS-CoV-2 variants of concern (VOCs), leading to increased transmissibility and immune evasion ^7,8^. For example, the N501Y mutation, present in the Alpha and Omicron variants, enhances the binding affinity to hACE2, while the E484K and L452R, found in Beta, Gamma, and Delta, reduce the neutralization efficiency of therapeutic Abs ^9^. These mutations pose a persistent threat to the clinical efficacy of current therapeutic treatments.

Because viral entry begins with this molecular recognition step, the biophysical properties of the RBD/hACE2 interaction under mechanical stress are key to understanding infectivity and immune escape. Traditional affinity and neutralization data provide thermodynamic insights but do not fully capture the mechanical properties, which may be relevant in physiological and therapeutic contexts^10^. During infection, forces such as fluid shear stress, tissue deformation, and intracellular crowding act on viral and host components ^11^. Therefore, the mechanical stability of the RBD in complex with hACE2 or neutralizing agents has direct implications for viral attachment, immune evasion, and therapeutic resistance.

Recent studies using single-molecule force spectroscopy (SMFS) and molecular dynamics (MD) simulations have shown that the mechanical stability of RBD/hACE2 complexes varies across SARS-CoV-2 variants ^7^. Notably, variants such as Alpha, Beta, and Omicron exhibit increased force resistance at the RBD/hACE2 interface, consistent with enhanced binding affinity and transmissibility. However, while the nanomechanics of hACE2 complexes have been extensively studied, the nanomechanical properties of SARS-CoV-2 RBD binding neutralizing Abs and Nb remain largely unexplored^12–15^.

Abs and Nbs differ in their size, structure, and binding mechanisms. Conventional Abs are bivalent ~150 kDa molecules composed of heavy (H) and light (L) chains, while Nbs are an order of magnitude smaller single-domain proteins derived from camelid heavy-chain-only Abs ^16,17^. These structural differences influence not only epitope accessibility but also the distribution and transmission of mechanical forces during complex dissociation. Understanding how these biomolecules behave under force is essential for designing more robust therapeutics. Particularly, Nbs have attracted attention for their favorable biophysical properties, including high solubility, thermal stability, and the ability to bind cryptic epitopes on viral proteins ^17^. Several Nbs have shown potent neutralization against SARS-CoV-2. Among these, the llama-derived Nb H11-H4 binds a conserved epitope on the RBD with nanomolar affinity and has been shown to block viral entry in vitro^18^.

Although SMFS provides valuable insights into rupture forces of Abs or Nb-antigen complexes ^7,19^, it cannot fully resolve the dissociation pathway under mechanical load. MD simulations offer a complementary perspective, providing atomistic or coarse-grained (CG) resolution^20^ of protein motions under force^21^. Steered molecular dynamics (SMD) simulation has been widely applied to study protein-protein dissociation ^21–23^. However, all-atom (AA) SMD simulations are computationally expensive and often require high pulling speeds that may overestimate rupture forces and distort the unbinding pathways^24^.

To address these limitations, we employed the GōMartini 3 ^25^, a methodology that combines the Martini 3 force field ^26^ with a structure-based model ^27^. In this framework, contacts are defined based on an optimised protein contact map that considers the overlap (OV) of the van der Waals radii and the repulsive contacts of structural units (rCSU) between heavy atoms ^28^. GōMartini 3 enables accurate mechanical characterization of large biomolecular complexes at reduced computational cost while allowing for slower and more realistic pulling velocities ^12,25^. Unlike traditional CG models that rely on harmonic restraints, GōMartini 3 preserves the tertiary and quaternary structures of proteins and permits large-scale conformational changes^25^.

In this study, we performed a comparative nanomechanical analysis of six RBD complexes comprising three Abs and three Nbs, each binding a distinct region of the RBD (Figure 1). We selected systems with available experimental structures and modeled two SARS-CoV-2 Omicron variants, BA.4 and JN.1, to assess the effects of mutations on mechanical behavior. Each complex was first relaxed using AA-MD simulations and then subjected to CG-SMD simulations using GōMartini 3. In addition to intact Abs, we designed single-chain systems in which either the H or L chain was bound alone to the RBD, enabling us to probe the individual contributions of each chain to force transmission. By analyzing the rupture of contacts under force, we compared the nanomechanical resilience of Ab- and Nb-bound RBD complexes, and dissected how cooperative versus individual chain contributions shape their response. Our findings reveal how structural architecture and binding orientation govern the nanomechanical stability of these immune complexes, offering insights for the rational design of therapeutics with enhanced mechanical robustness.

**Figure 1.**
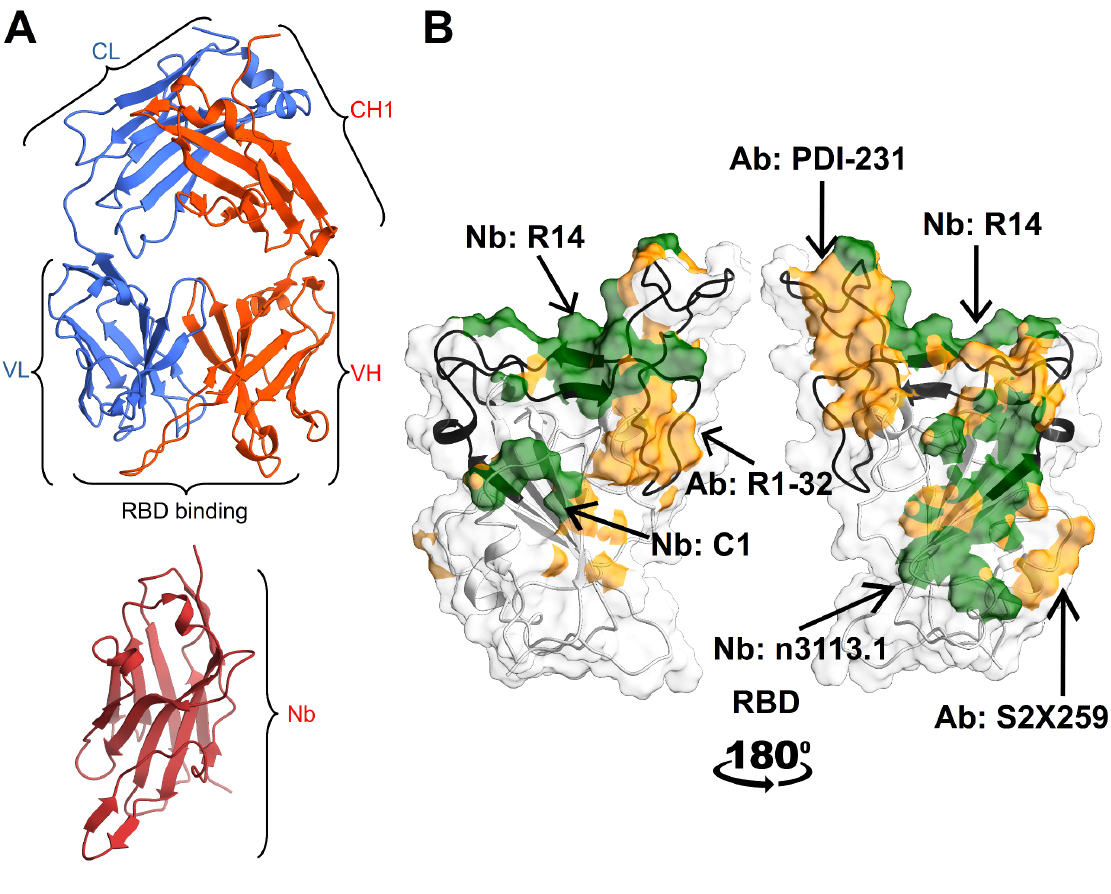
Structural overview of Ab and Nb recognition of the SARS-CoV-2 RBD. **A)** Domain organization of a conventional Ab (top) comprising the L chain (variable region, VL, and constant region, CL, shown in blue) and the H chain (variable region, VH, and first constant domain, CH1, shown in orange), and the bottom shows the single-domain variable region of a heavy-chain-only Ab (Nb, shown in red). **B)** Representative binding footprints of Abs (PDI-231, R1-32, S2X259) and Nbs (R14, C1, n3113.1) on the RBD (cartoon in black, surface representation in gray). Epitope regions targeted by each Ab and Nb are highlighted on the RBD surface in orange and green. The two panels show the RBD rotated by 180° to visualize binding epitopes from opposite orientations. The RBM region is highlighted in black color.

## 2 Methods

### 2.1 Molecular modeling

We first generated complete structural models of Ab and Nb complexes with the SARS-CoV-2 RBD, which served as an input for subsequent relaxation and nanomechanical analysis. A total of six representative complexes were selected, targeting three distinct epitopic regions of the RBD. These included three Abs, PDI-231 (PDB ID: 7MZN)^29^, S2X259 (PDB ID: 7M7W)^30^, and R1-32 (PDB ID: 7YDI)^31^, and three Nbs, R14 (PDB ID: 7WD1)^32^, C1 (PDB ID: 7OAP)^33^, and n3113.1 (PDB ID: 7VNE)^34^. Missing residues in 7M7W (residues 141-147) were modeled using Modeller v10.5^35^. In the 7YDI structure, hACE2 was removed, while for 7VNE, only the RBD/Nb portion was retained. To avoid artificial termini charges, the N- and C-termini of the RBD, H and L chains, and Nb (where needed) were capped with acetyl and methylamide groups, respectively. Additionally, BA.4 and JN.1 SARS-CoV-2 Omicron variants were modeled using the WT RBD as a template, incorporating the corresponding mutations via Modeller while preserving the original Ab or Nb binding geometry. Table 1 summarizes the structural and binding features of Ab and Nb targeting the WT RBD. A full list of protein contacts for all complexes is provided in Supplementary Tables S2-S7.

**Table 1.**
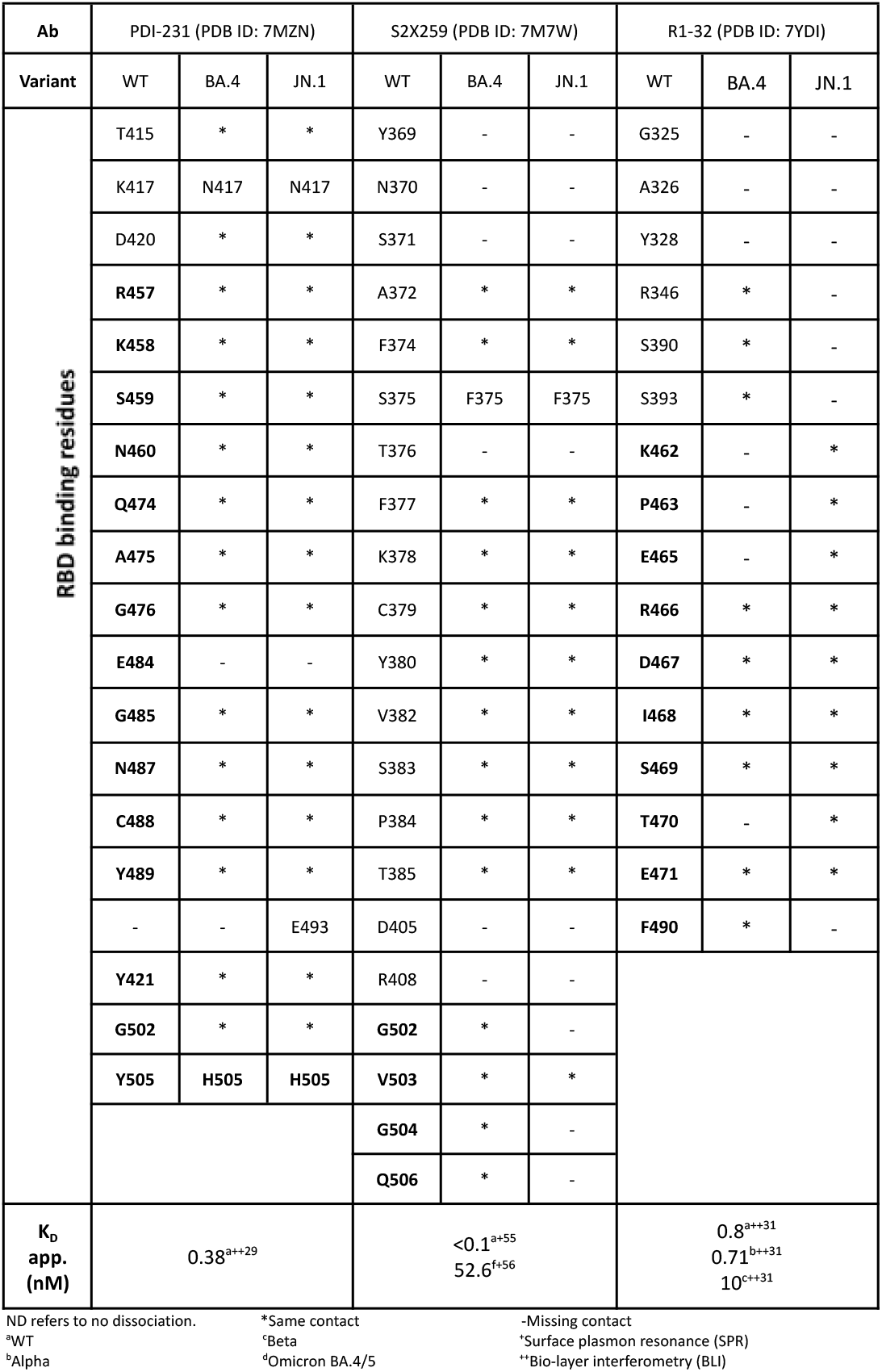
Structural and binding parameters of the selected Abs targeting the SARS-CoV-2 RBD. For each protein case, the name, type, PDB ID, and RBD residues making contacts with the WT are listed. Experimental dissociation constant (K_D_) parameters considering different variants are reported. Residues in bold are part of the RBM region.

### 2.2 AA-MD simulations

AA-MD simulations were then performed to relax each complex and to obtain equilibrated structures suitable for CG modeling within the GōMartini 3 framework. These simulations were carried out using the AMBER 24 software package with the ff19SB force field ^36,37^. Simulations were run with the GPU-accelerated pmemd.cuda engine^38^. Initial energy minimization in a vacuum was performed for each complex to eliminate steric clashes, particularly around mutated residues introduced during modeling. The protonation state of titrable residues was assigned at pH 7.4 using PDBFixer^39^.

Each minimized complex was solvated in a dodecahedral box of explicit water molecules using the four-site OPC water model^40^, with a minimum distance of 10 Å between any protein atom and the box edge. System neutrality was achieved by adding Cl-counterions, with the number of ions adjusted based on the net charge of each system. The number of ions added ranged from 2 to 12, depending on the SARS-CoV-2 variant (i.e. WT, BA.4 and JN.1).

Following solvation, systems underwent 5,000 steps of energy minimization using the steepest descent algorithm to resolve unfavorable contacts and to allow for solvent relaxation. Temperature equilibration was performed under constant volume (NVT) conditions by incrementally increasing the temperature from 150 K to 310 K in five 200-ps steps. During this phase, harmonic position restraints were applied to protein heavy atoms, with spring constants progressively reduced from 5 to 1 kcal mol^−1^ Å^−2^ to permit gradual relaxation of the protein backbone and side chains. This was followed by a 1 ns unrestrained equilibration at 310 K under constant pressure (NPT) conditions.

Production simulations were conducted in the NPT ensemble using periodic boundary conditions. Long-range electrostatics were treated with the particle mesh Ewald method, employing a grid spacing of 1 Å. Short-range interactions were modeled using a Lennard-Jones (LJ) potential with a 9 Å cutoff ^41,42^. Temperature was regulated using Langevin dynamics with a collision frequency of 4 ps^−1 43^, while pressure was maintained at 1 bar using the Monte Carlo barostat with a relaxation time of 2 ps^44^. Covalent bonds involving hydrogen atoms were constrained using the SHAKE algorithm^45^, and hydrogen mass repartitioning was applied using ParmEd^46^, allowing the use of a 4 fs integration time step^47^. Each system was simulated for 500 ns. All complexes remained structurally stable throughout the simulations.

### 2.3 Contact map determination

To identify high-frequency contacts at the interface of each complex from AA-MD simulations, we analyzed the atomistic MD trajectories of Ab or Nb/RBD systems. Each system was simulated for 500 nanoseconds, and contacts were calculated every 1 nanosecond using a Python script (https://github.com/GoMartini3-tools/ContactFreq). This script uses the executable version of the Go Contact Map from the server (http://pomalab.ippt.pan.pl/GoContactMap/) which is also available at https://zenodo.org/records/3817447 and https://github.com/Martini-Force-Field-Initiative/GoMartini/tree/main/ContactMapGenerator.

We considered both the van der Waals (vdW) radii overlap (OV) and repulsive chemical structural units (rCSU) contact map methods ^28,48^. The OV contact map method is a purely geometric criterion that identifies interactions based on the spatial overlap of vdW spheres centered on heavy atoms. To incorporate attractive contributions, each vdW radius is scaled by a factor of 1.24. A contact is defined when spheres from two residues, separated by at least four positions in the sequence, overlap. This approach has been widely used to identify stabilizing interactions relevant for folding and mechanical unfolding of protein domains ^12,21,23,27,48^. The rCSU method complements the OV approach by integrating chemical specificity and electrostatics. It considers both attractive and repulsive interactions and classifies residues as hydrophobic, hydrophilic, aromatic, or ionic. A contact is considered valid when the number of attractive interactions exceeds the number of repulsive ones. Each atom is represented as a sphere, and its surface is sampled using a Fibonacci grid, which ensures a uniform and unbiased distribution of points. Only high-frequency contacts, defined as those present in more than 70 percent of the frames, were retained. The frame containing the largest number of contacts was selected for CG modeling using the GōMartini 3 framework (Tables S2-S7).

### 2.4 CG-MD simulations

CG models for Ab and Nb complexes were prepared using the Martini 3 force field^26^ and the martinize2 tool^49^. Secondary structure definitions were assigned using DSSP v3.0 ^50^. We applied the GōMartini 3 approach ^25,27^, which incorporates a Lennard-Jones potential between virtual sites derived from the high-frequency contact map ^51^. The depth of the LJ potential for contact pairs was set to 15 kJ mol^−1 12,21^. The CG structures were initially minimized in a vacuum during 5000 steps using the steepest descent algorithm. Subsequently, the complexes were solvated in a 2 × 2 × 2 nm^3^ box with the standard Martini water model. Sodium and chloride ions were added to neutralize each system and to achieve a physiological strength of 0.15 M. Positional restraints were imposed on the BB beads of each protein to prevent drifting during the equilibration phases. Both the NVT and NPT equilibrations, along with the production phase, utilized the V-rescale thermostat ^52^. The temperature coupling time constant was set at 1.0 ps for both protein and solvent components of the system, keeping the temperature at 300 K. The NVT equilibration was run for 2 ns, with an integration time of 20 fs. During the NPT equilibration and production, an isotropic pressure coupling was used with a compressibility set at 10^−4^ bar^−1^ and 1 bar pressure. The NPT equilibration was run for 5 ns using the C-rescale barostat^53^ with a pressure coupling time constant of 12 ps and an integration time of 10 fs. For the production phase, the C-rescale barostat was used, with a pressure coupling time constant of 15 ps. The cutoff distances for Coulomb and van der Waals (vdW) interactions were set at 1.2 nm across all equilibration and production phases. The equilibrium simulations were conducted over 2 microseconds, with a time step of 20 fs. A total of 5 independent replicas were conducted for each system using GROMACS 2023.5 ^54^.

We monitored interfacial contact frequency over 2 μs equilibrium simulations (Figures S16–S19). All RBD/Ab and RBD/Nb complexes preserved their characteristic binding interfaces, with contacts maintained with high frequency throughout the trajectories. While some variant-specific contacts showed moderate fluctuations, the dominant interaction networks remained stable across replicas. These results indicate that the GōMartini parametrization reliably sustains the binding modes of the complexes under equilibrium conditions, providing robust starting points for the nanomechanical pulling simulations.

### 2.5 CG-SMD simulations

Systems were prepared as stated in the previous subsection. All CG systems were initially energy-minimized in a vacuum for 5,000 steps using the steepest descent algorithm. To accommodate the pulling geometry, the complexes involving conventional Abs were solvated in a 10 × 10 × 100 nm^3^ water box, while the RBD/Nb complexes were solvated in a 10 × 10 × 80 nm^3^ box. Minimization and equilibration cycles were the same as above. Production simulations used the same barostat with a coupling time of 8 ps and a 20 fs integration time step. The cutoff distance for both Coulombic and van der Waals interactions was set to 1.2 nm throughout all stages of the CG simulations.

SMD simulations were carried out for 2.5-3.0 μs per replica. For each complex, directional constraints were applied to mimic mechanical dissociation. Specifically, the positions of the heavy atoms of the last three residues at the C-terminus of the RBD were restrained along the pulling direction (i.e. the z-axis). Simultaneously, the coordinates of the last three residues of the Nb or the Ab’s H or L chain were fixed in the x- and y-directions. The pulling force was applied to the center of mass (COM) of the selected terminus using a constant velocity of 1 × 10^−5^ nm ps^−1^ and a harmonic spring constant of 37.6 kJ mol^−1^ nm^−2^. For conventional Abs, two separate pulling sets were defined: i) set 1, pulling was carried out from the H chain, and ii) set 2, pulling was performed out from the L chain. A total of 50 independent replicas were performed for each configuration using GROMACS 2023.5^54^. In addition to these full-complex simulations, we constructed single-chain systems in which either the H or the L chain was removed prior to simulation. These reduced systems were subjected to the same pulling protocol, enabling us to isolate and evaluate the individual contributions of each chain to the overall mechanical response.

To evaluate the dynamic stability of protein–protein interfaces during CG pulling simulations, we systematically monitored the rupture of contacts throughout each trajectory. The analysis was based on an effective representation of atoms as enlarged vdW spheres, which better approximate the spatial extent of atoms by accounting for the influence of their electron clouds. Contact pairs were defined according to the interaction list specified in the GōMartini framework. For each contact, a specific σ value was assigned, corresponding to the distance at which the LJ potential equals zero. The position of the potential energy minimum (R_min_) was then calculated as σ multiplied by 2^⅙^. During the simulations, we computed the distance between the BB beads of each contact pair at every frame and compared it to a scaled threshold, set to 1.3 times R_min_, following previous GōMartini studies ^12^. A contact was considered intact if the BB-BB distance was less than or equal to this threshold; otherwise, it was classified as broken.

## 3 Results & discussion

### 3.1. Molecular Basis of Ab–RBD Interactions

To investigate the molecular determinants of stability in the RBD–PDI-231 complexes, we analyzed the chemical nature of the interfacial contacts between the Ab and the SARS-CoV-2 RBD variants. The H chain comprises a variable domain (VH, residues 1–124) and a constant domain (CH1, residues 133–228), while the L chain consists of a variable domain (VL, residues 1–112) and a constant domain (CL, residues 121–217) ^29^. The interface is mainly formed by VH and CH1 from the H chain and the VL domain from the L chain. Contact analysis revealed a network of persistent hydrophobic, polar, and aromatic interactions across variants. K378 from the RBD interacts with Y102, Y103, and W105 from VH, suggesting cation–π interactions, while Y380 forms hydrophobic contacts with N101 and Y102. CH1 contributes polar contacts with RBD residues D405 and G502 via CL residues. A conserved RBD segment (residues 515–529) engages both VH and VL domains. In the WT complex, residues such as F490, Y489, and A475 form a hydrophobic core, preserved in BA.4 and JN.1 despite surrounding mutations. In contrast, ionic interactions observed in the WT complex, including K458–E50 and R457–D52, are lost in Omicron variants due to mutations such as K417N and E484A. Despite these changes, CL-mediated interactions (e.g., Y103 and Y52 with Y421 and G485) remain conserved, highlighting the stabilizing role of the L chain (Table).

Analysis of the WT/S2X259 complex revealed a predominantly hydrophobic interface, particularly centered around VH residues (residues 1–124) such as A375, C379, S375, and F377 forming frequent contacts with RBD residues Y505, F456, and G476. These hydrophobic interactions are mainly localized within the complementarity-determining regions (CDRs), particularly CDR1_H (residues 25–33), CDR2_H (residues 52–58), and CDR3_H (residues 100–112), and form a compact, structurally resilient core ^30^. In contrast to PDI-231, S2X259 shows fewer ionic interactions. Notably, the K378–Y505 and K378–G476 ionic pairs observed in the WT complex are disrupted in the BA.4 and JN.1 variants due to mutations, reflecting diminished electrostatic complementarity. Despite this, the BA.4 variant preserves many of the hydrophobic interactions, such as S375–Y505 and C379–F456, maintaining a similar interfacial architecture. In JN.1, the number of interfacial contacts decreases markedly, especially those involving the CL domain (residues 121–217), where all contacts are lost. Nevertheless, conserved interactions remain between VH and RBD, such as A375–Y505 and F377–F456, supporting the idea that the VH domain of S2X259 maintains a degree of interfacial stability despite RBD mutations. Overall, the data suggest that S2X259 relies primarily on hydrophobic contacts mediated by VH for RBD recognition, while the contribution from CL is significantly reduced in the Omicron variants (Tables 1 and S2).

The H chain of R1-32 comprises a variable domain (VH, residues 1–122) followed by a constant domain (CH1, residues 131–228), while the L chain contains a variable domain (VL, residues 1–110) and a constant domain (CL, residues 119–214) ^31^. In the WT complex, the interface is dominated by hydrophobic contacts established by the VH domain, particularly involving residues in CDR1 and CDR2 (e.g., E484–Y102, F486–Y52, F486–Y103, and Y489–Y52), suggesting the formation of a robust hydrophobic core that anchors the Ab to the RBD. Polar interactions such as D467–Y102 and E465–Y105 further stabilize this interface through hydrogen bonding and electrostatic complementarity. The CH1 domain contributes fewer interactions but remains involved through residues like S409 and T470. Comparison with the BA.4 and JN.1 complexes reveals both conserved and disrupted interfacial interactions. Notably, key hydrophobic contacts such as E484–Y102 and F486–Y103 are retained in both variants. However, BA.4 and JN.1 lose several contacts mediated by CH1 and CL residues, including those involving S409, T470, and S390. The L chain in particular contributes fewer contacts in the Omicron variants, with interactions like S390–K528 observed only in WT. Electrostatic interactions, such as K356–S391 and N354–S391 in BA.4, and N354–S391 in JN.1, emerge as variant-specific features, potentially compensating for the loss of hydrophobic contacts. Overall, the VH domain remains the primary contributor to binding across all variants, with variant-specific remodeling occurring mainly in contacts involving CH1 and CL.

### 3.2 Molecular Basis of Nb–RBD Interactions

R14 engages the RBM through a compact interface dominated by hydrophobic and aromatic contacts, with key contributions from F490 (interacting with residues Y104, Y109, T102 from the Nb) and Y449 (contacting residues A101, Y103, Y31 from the Nb) ^32^. Additional polar stabilization is provided by Q493–T102. In the WT complex, multiple hydrophobic contacts involve F486, but these are lost in BA.4 and JN.1, reducing the hydrophobic packing at the interface. JN.1 exhibits the fewest total contacts among the three complexes, indicating a more disrupted interface. Compared with PDI-231, which combines hydrophobic, polar, and several ionic interactions (e.g., K458–E50 and R457–D52 in WT), R14 relies less on salt bridges and more on a localized hydrophobic/aromatic core. This contact profile may contribute to its lower affinity for Omicron BA.4 (K_D_ = 43.3 nM) relative to WT, where no dissociation was detected.

C1/RBD binding surface partially overlaps the S2X259 epitope^33^. In the WT complex, recurrent interactions include K378–A102/D105/F101/G103, F377–G103, G381–F31, and hydrophobic pairs Y380–F101 and V382–F31, supported by a polar cluster around S383 (with S54 and T106) and R408–S109. Many of these contacts persist in BA.4, with additional ionic pairs involving K378–D105 and K378–R104. In JN.1, the interface remains hydrophobically anchored (Y380–F101, V382–F31). Compared with S2X259, which presents a hydrophobic core involving F486, Y489, and A475, it loses several polar or ionic contacts in BA.4 and especially JN.1. C1 retains its key hydrophobic pairs and introduces alternative polar or ionic interactions around K378.

n3113.1 forms a compact interface centered on a hydrophobic pocket that involves A348/A352 of the RBD contacting A100 of the Nb, together with I468–Y32 and the conserved Y449–W47 pair^34^. These interactions are complemented by a polar network involving R346–D106 and residues around N354/T345, whose specific contact pairs shift between variants. In WT, N354–S101 and T345–S103 are observed, whereas BA.4 retains T345–S103 but lacks N354 contacts. In JN.1, N354 interacts with S101 or T104, and a new T356–T104 contact emerges. Total contacts drop in BA.4 (n=17) but recover in JN.1 (n=23), with added polar contacts such as N354–S101/T104 and T356–T104, plus D450–W47. In comparison, R1-32 presents a broader interface dominated by VH contacts to E484, F486, Y489, and related RBM residues, with additional contributions from CH1 and occasional CL contacts. Across BA.4 and JN.1, several constant-domain and light-chain interactions are lost, and the interface relies increasingly on the VH hydrophobic core.

Overall, the Abs and Nbs examined displayed distinct strategies for RBD recognition, ranging from broad, mixed hydrophobic–polar interfaces (PDI-231, R1-32) to more localized hydrophobic/aromatic cores (R14, C1, n3113.1). Omicron BA.4 and JN.1 disrupt several ionic and peripheral contacts, particularly in constant-domain and light-chain regions, while core VH or Nb–RBD hydrophobic contacts are generally preserved. R14 and S2X259 showed marked contact loss in JN.1, consistent with reduced affinity, whereas C1 and n3113.1 compensate through alternative polar or ionic interactions. These patterns highlight both the vulnerability of certain epitopes to variant mutations and the capacity of others to adapt via contact reorganization.

### 3.3 Nanomechanics of Ab/RBD complexes

The binding affinity of Abs has been extensively characterized ^57^. More recently, we demonstrate the inherent mechanical stability present in potent neutralizing Nbs across different SARS-CoV-2 variants ^12^, but the relative contribution of the H and L chains to mechanical stability remains poorly understood. To address this gap, we employed SMD simulations to probe the Ab response under force by applying constant-velocity pulling either from the C-terminus of the H chain or from the L chain in independent simulations. This setup allowed us to monitor force-displacement profiles, identify dissociation pathways, and assess the stabilizing role of inter-chain contacts during mechanical stress.

#### 3.3.1 PDI-231/RBD complex

Both PDI-231 and R14 are representative hACE2-competing binders that neutralize SARS-CoV-2 by preventing the spike protein from interacting with the hACE2 receptor. PDI-231 is a monoclonal Ab that binds directly to the RBM of the RBD, overlapping with the hACE2 binding site ^29^. It exhibits a K_D_ of 0.38 nM against the WT strain and retains potent neutralizing activity against multiple variants, including those carrying key mutations such as N501Y, E484K, and K417N ^29^. However, due to its epitope positioning, PDI-231 can only engage the RBD when it adopts the up conformation ^29^. We first analyzed the mechanical dissociation behavior of the PDI-231/RBD WT complex using 50 independent CG-SMD simulations (Figure 2A). When a constant pulling velocity was applied to the H chain, two distinct dissociation behaviors were identified. In the predominant case, the force–displacement profile displayed a single pronounced peak (Figure 2A), consistent with rigid-body dissociation, in which both Ab chains (H in red and L in blue) and the RBD (black) remained folded from the point of maximum force until full dissociation. In approximately 28% of SMD trajectories, partial unfolding sampled by the reduction of contacts in the C-terminal region of the H chain (residues V207–D226) was observed before dissociation, whereas the RBD remained structurally stable as it maintained most of all initial contacts across pulling (Figure S1A). Across all CG-SMD trajectories, the H chain was consistently the last component to detach from the RBD.

**Figure 2.**
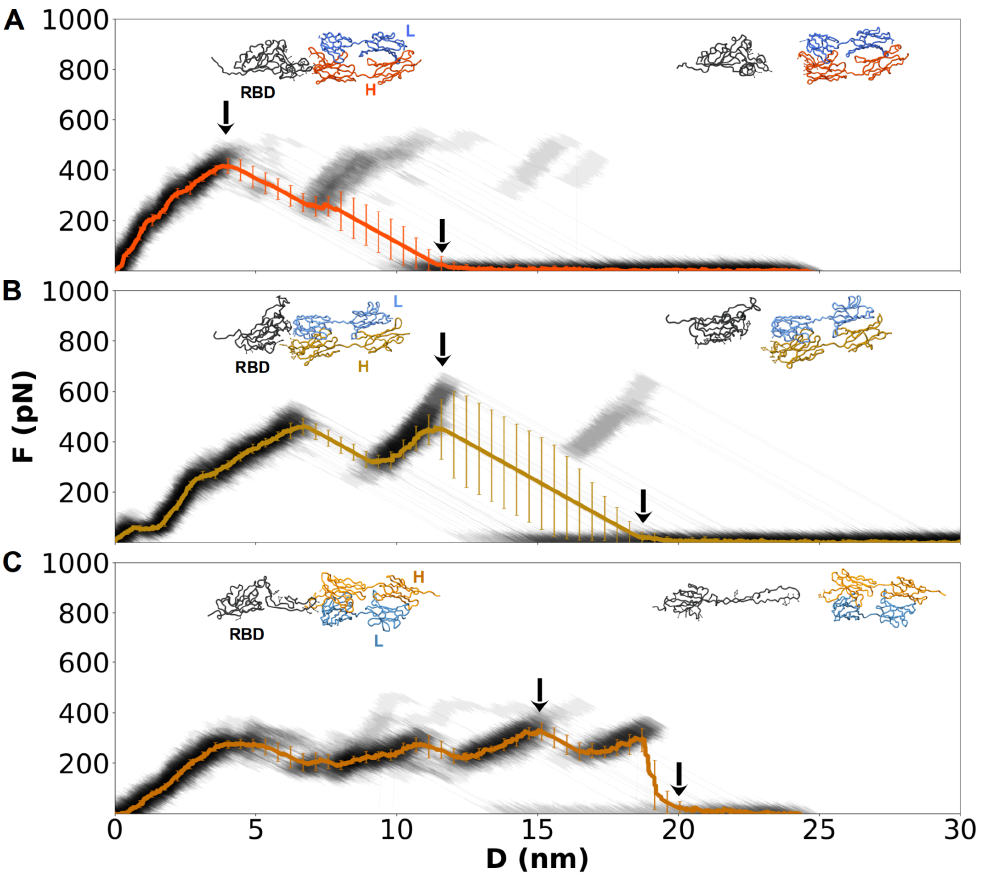
Mechanical response of Ab/RBD WT complexes under constant-velocity pulling from the H chain. Force–displacement profiles are shown for (A) PDI-231, (B) S2X259, and (C) R1-32 (gray traces, mean in color). Insets display representative structures at the maximum force (F_max_, left arrow) and after dissociation (right arrow). Error bars denote standard deviations across 50 independent trajectories.

When SMD pulling was applied to the L chain in the WT complex (Figure S5A), dissociation also proceeded predominantly via a rigid-body mechanism. Dissociation from the L chain was often accompanied by unfolding of RBD residues A522–H530 (Figure S1B). This reflects the H chain’s role as the main load-bearing element; it absorbs much of the applied force, limiting RBD deformation. In contrast, the weaker L chain transmits force more directly into the RBD, promoting local destabilization and unfolding within the A522–H530 segment.

Figure S12 and Table S2 show a comparative analysis of inter-chain contacts in the Ab PDI-231 bound to the three different SARS-CoV-2 RBD variants. In the PDI-231/RBD complex with the WT variant, pulling from the H chain led to the loss of many contacts between the H and L chains. The most prominent contact to fail under force was S114–S141, a constant region interaction that consistently broke early during pulling. Additional contacts, including F118–L133 and T164–F175, also located within the constant domains, showed reduced stability. The loss of pairs such as S162-P176 and V163-P176 further highlights that the mid-to-C-terminal segment of CH1 detaches readily from its CL partner. Pulling from the L chain produced a similar outcome: the S114–S141 contact again failed first, indicating that this region was mechanically vulnerable regardless of the pulled chain.

For the RBD/PDI-231 complex with the BA.4 variant, pulling from the H chain also led primarily to rigid-body dissociation, with the L chain consistently detaching before the H chain (Figure S8A). Unfolding of the C-terminal region of the H chain (residues I204–D226) preceded rupture and was observed in some trajectories. In rare cases, the residues K215–D226 from the constant region of the H chain underwent limited unfolding before dissociation, while the L chain remained folded (Figures S2A and S8A). When pulling from the L chain in the BA.4 complex, most dissociation events followed a rigid-body profile, with the L chain detaching first (Figure S6A). Partial unfolding of the L chain’s C-terminal region (residues E195–C214) occurred before detachment in 4% of the trajectories (Figure S2C). When pulling from the H chain, we observed that seven pairs of contacts were mechanically weakened in comparison with the WT (Table S9, Figure S12). These pairs of contacts (T192-N137, V178-Q160, P176-S162, P176-V163, F175-S162, F175-V163, F175-T164) are located in the CH1 and CL domains, respectively. Contacts A146-F116, P135-F118, and A134-F118 also presented a reduced mechanostability to a lower extent. Interestingly, when pulling from the L chain, all inter-chain contacts remained present, which is consistent with the rigid-body dissociation pattern observed for these simulations (Figure S12).

For the JN.1 variant, pulling from the H chain revealed two dissociation modes. The dominant profile involved a single force peak and corresponded to rigid-body dissociation. In these simulations, partial unfolding of the constant regions of both chains in the Ab was observed (Figure 3A). The L chain consistently detached first, followed by the H chain. When CG-SMD pulling was applied to the L chain, a similar rigid-body dissociation was observed, with the H chain being the last to separate. These trajectories were also characterized by a single dissociation peak. The L chain initially broke contacts at the constant region and then dissociated from the RBD. The H chain remained attached during this phase. In several trajectories, the L chain unfolded between residues Q200–E216 before its dissociation. The L133-F118 contact was the most affected mechanically when pulling from either the H chain or the L chain. The same set of residues was mechanically destabilised when pulled, regardless of the chain (Figures 3A, S3A/B and S12). The most affected contacts when pulling the H chain were F175-S162, P176-V163, F175-T164, and A146-F116, whereas pulling from the L chain affected P132-S121, F131-S121, F131-E123, and F131-Q124.

**Figure 3.**
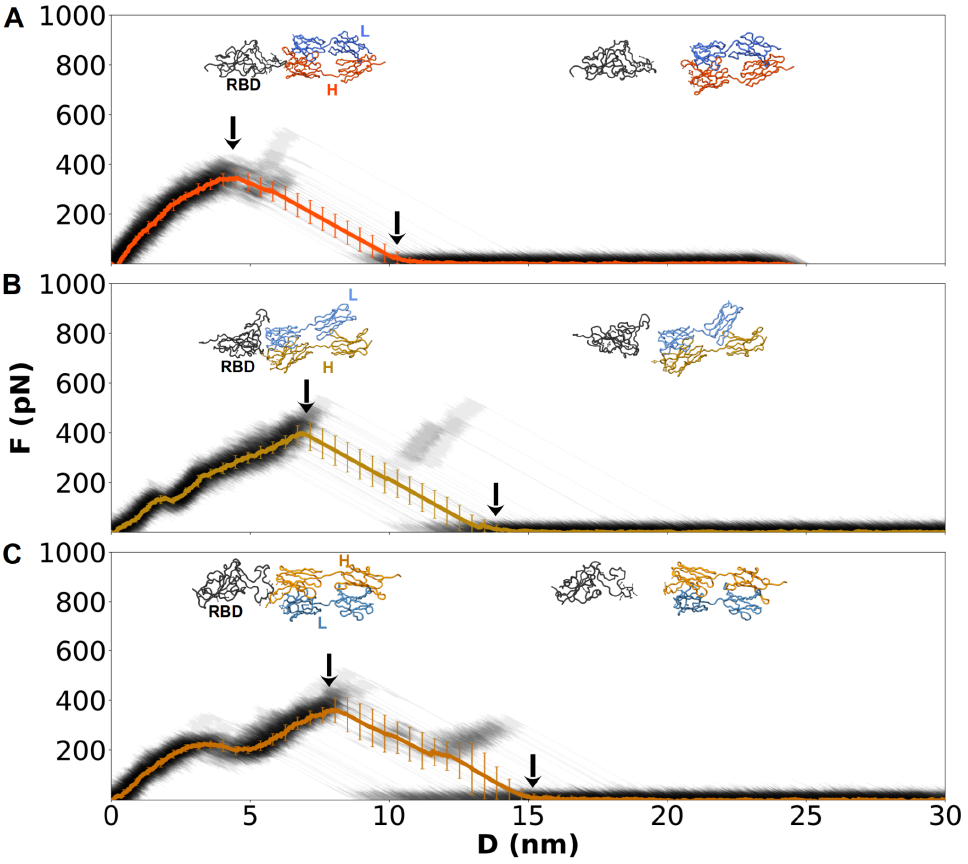
Mechanical response of Ab/RBD JN.1 complexes under constant-velocity pulling from the H chain. Same as Figure 2.

#### 3.3.2 S2X259/RBD complex

S2X259 is a broadly neutralizing Ab with exceptional affinity for the SARS-CoV-2 WT strain, displaying a K_D_ below 0.1 nM ^30,34^. It retains potent neutralizing activity against multiple variants of concern, including Omicron, with K_D_ ranging from 1 to 86 nM, and is also effective against several zoonotic sarbecoviruses ^55,58^. S2X259 binds a cryptic and structurally conserved RBD epitope composed of residues Y369–K386, G404–A411, and P499–Y508. This epitope is only exposed in the open state of the spike protein. Although it does not directly contact the RBM, S2X259 prevents hACE2 engagement through steric hindrance, physically obstructing hACE2 binding to the RBD ^55^ (Table S9). In this Ab, the VH domain spans residues Q1-127A, followed by CH1 from S133-S228. The VL domain includes residues Q1-V112, while CL extends from S121-E217.

In the WT, when pulling from the H chain, most trajectories exhibited a single dissociation peak (Figure 2B). In these cases, the RBD underwent unfolding of residues P521–K529. This mechanical destabilization was consistently followed by the dissociation of the H chain from the RBD. Subsequently, the L chain detached, completing the separation of the Ab from the RBD. In some trajectories, however, the same region of the RBD unfolded, but residues N208–S226 of the H chain also showed signs of unfolding before dissociation (Figures S1C and S13). In these cases, the H chain still dissociated first, and the constant region of the Ab occasionally lost inter-chain contacts before complete unbinding.

For the BA.4 variant, pulling from the H chain consistently resulted in the unfolding of RBD residues A520–S529, after which the H chain dissociated first, followed by the L chain (Figure S6). In most trajectories, no additional unfolding event was observed along the Ab or in other regions of the RBD. However, in 10% of the trajectories, the H chain unfolded in residues V122–T146 and H211–S226, indicating a destabilization of its VH and CH1 regions prior to dissociation.

When pulling from the L chain, the RBD consistently unfolds in the A520–S529 region (Figure S5B). This was followed by progressive unfolding of the L chain, first across residues Q200–E216, then extending to residues L112–K135. After these unfolding events, the L chain was dissociated completely from the RBD, and all contacts were lost with the H chain. Notably, this chain remained bound to the RBD in these cases, suggesting an asymmetric mechanical response. In this case, only the L chain unfolded and dissociated.

The mechanical response for the JN.1 variant was more heterogeneous (Figure 3B). When pulling from the H chain, some trajectories showed loss of contacts within the constant regions, followed by the simultaneous detachment of both chains from the RBD. These cases did not involve unfolding in the Ab chains, and the only structural perturbation in the RBD was localized to residues L516–K528. In other trajectories, the H chain exhibited unfolding in the N210–S226 region, while the L chain dissociated first, followed later by the H chain.

When pulling from the L chain, two distinct behaviors were observed. In some cases, a rigid-body dissociation occurred, where this chain detached first without any noticeable unfolding in the protein complex (Figure S6B). In other trajectories, the L chain unfolded between residues T202–E216, or even more broadly from S198 to E216, before dissociating from the RBD. These events were followed by loss of some contacts with the VL region, and eventually, the H chain also separated, completing the dissociation event.

Across all three variants, the RBD consistently unfolded near the C-terminal region (residues A516–S529) in response to mechanical forces. The order of dissociation was typically defined by the pulled chain, though in the JN.1 variant, simultaneous detachment or variable dissociation orders were occasionally observed. Ab unfolding was relatively rare, highlighting the high degree of mechanical stability present and only limited to specific regions: the H chain (V122–T146 and 208–226) or the L chain (Q200–E216 and D112–L135), depending on the variant and pulling direction (Figures S1-3 and Table S3). These unfolding events were more pronounced in BA.4, particularly when pulling from the H chain. Meanwhile, the JN.1 variant tended to show less extensive unfolding and, in some cases, more rigid-body-like dissociation. This possibly indicates a shift in mechanical stability and inter-chain interactions in this variant.

#### 3.3.3 R1-32/RBD complex

R1-32 is a monoclonal Ab that binds a cryptic, conserved epitope located beneath the RBM of the SARS-CoV-2 RBD ^31^. This site becomes fully accessible only when the RBD adopts the open (up) state. This Ab does not compete with hACE2 and does not act through steric hindrance (Table S10). Instead, it promotes RBD opening and destabilizes the spike protein, potentially driving the complex into a non-functional conformation ^31^. For the WT complex, pulling from the H chain showed dissociation profiles with a sawtooth pattern (Figure 2C). The RBD consistently unfolded in the S438–P507 region, while the Ab chains remained structurally intact (Figures S1E/F, S14 and Table S4). Dissociation proceeded with the L chain detaching first from the RBD, followed by rupture of contacts between the constant regions of the Ab and eventual dissociation of the H chain. Pulling from the L chain yielded similar profiles: the RBD unfolded in the same S438–P507 region, the L chain dissociated first, and the H chain was the last to break protein contacts. No unfolding events were observed in either Ab chain during these simulations.

The unfolding of the S438–P507 region could be caused by the intrinsic mechanical weak points of the RBD identified in prior pulling simulations ^48^. These simulations revealed that the peak rupture force (F_max_) in the RBD is associated with the loss of contacts involving residues A348, S349, V350, and Y351 in the β1 region (Figure S15), as well as loop elements contacting V401, L452, Y451, Y453, R454, and L492 (β5 and β6 regions). Disruption of these contact clusters destabilizes the β-sheet core and the loop preceding the RBM, propagating to β5–β6 and facilitating β-strand separation. The proximity of S438–P507 to these mechanically labile contact regions explains why this segment reproducibly unfolds under force in the R1-32 complex.

For the BA.4 variant, pulling from the H chain produced dissociation events in which the L chain consistently detached first, followed by RBD unfolding (residues R454–P491) and then rigid-body dissociation of the Ab (Figure S6C). No unfolding was observed in either chain of the Ab. In 74% of the trajectories, RBD unfolding extended to residues S438–P507, again with the L chain detaching first. Pulling from the L chain yielded similar RBD unfolding in the S438–P507 region in some simulations (Figures S6C, S2E/F, S14). Additionally, the L chain showed partial unfolding in residues V199–T213 before dissociation. In all cases, the L chain dissociated first from the RBD.

In the JN.1 variant, pulling from the H chain resulted in dissociation trajectories dominated by RBD unfolding in residues S438–Y507, consistent with the WT response (Figures 3C and S6C). The L chain again detached first from the RBD, followed by the H chain. The Ab chains remained fully folded throughout. In some trajectories, unfolding was more localized, affecting residues R454–L491 within the RBD, but the overall mechanism remained the same. When pulling from the L chain, dissociation involved deformation of the RBD tip, followed by detachment of the L chain from the RBD. This was accompanied by disruption of the inter-chain contacts between L and H, leading to full separation of the L chain from the complex. Notably, the H chain remained bound to the RBD and did not unfold in any case.

#### 3.3.4 Cooperative role of heavy (H) and light (L) chains under mechanical forces

To dissect the contribution of individual chains to Ab mechanical stability, we carried out CG-SMD simulations in which either the H or L chain was bound alone to the RBD in the WT background. In all cases, the maximum force (F_max_) sustained by a single-chain construct was reduced compared to the two-chain Ab complex, underscoring the cooperative role of the complex architecture. The whole complex reached F_max_ values of about 365 pN for PDI-231, 550 pN for S2X259, and 408 pN for R1-32 (Figure 5). By contrast, H-only constructs supported forces of about 230–430 pN, whereas L-only systems reached an average value of 200 pN. These results indicate the non-additive dependency on the chain-type regarding the mechanical stability and that the two-chain AB complex provides synergistic stabilization via inter-chain contacts and the distinction of the mechanical load.

For PDI-231, the whole complex reached about 365 pN, while the H and L chains (in isolation) withstand about 250 pN and 200 pN, respectively (Figure 4). This shows that Ab complexes contribute meaningfully to mechanical processes. In the whole Ab system, H and L chains cooperate by distributing the mechanical force asymmetrically and delaying the Ab dissociation, increasing F_max_ beyond that reported by single chain studies. In S2X259, the whole Ab complex reached the highest F_max_ value at about 550 pN. Here, the H chain dominated, withstanding mechanical forces in the range of 430 pN, while the L chain remained in the range of 200 pN. The majority of stabilizing contacts are located at the H/RBD interface, giving rise to the asymmetry. The L chain provides a peripheral stabilization, detaching earlier and contributing less to the overall mechanical response. For R1-32, the whole Ab complex reached about 408 pN, nearly the same as the H chain (in isolation, 405 pN), while the L chain alone behaves mechanically similarly to previous cases. This indicates that force transmission is routed almost entirely through the H/RBD interface, with the L chain playing a non-negligible supportive role but not carrying the load.

**Figure 4.**
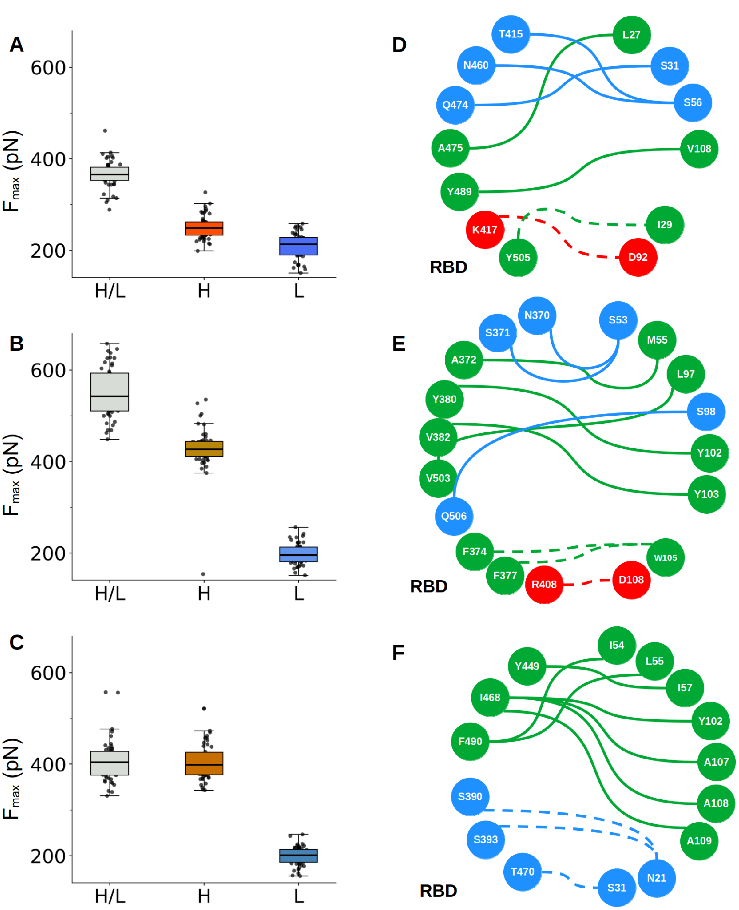
Mechanical stability and interfacial networks of Ab/RBD WT complexes. **A–C)** F_max_ values from CG-SMD simulations for PDI-231, S2X259, and R1-32, respectively. Boxplots compare the full Ab (H chain/L chain) with single-chain systems (H or L). Intact antibodies consistently sustain higher forces, reflecting cooperative stabilization between chains. **(D–F)** Network representation of contacts within the Ab/RBD interfaces of PDI-231, S2X259, and R1-32, respectively. Solid and dashed lines indicate H chain/RBD and L chain/RBD contact pairs, respectively. Colors denote interaction type: red for ionic, blue for polar, and green for nonpolar. Amino acid nodes are colored according to chemical properties.

**Figure 5.**
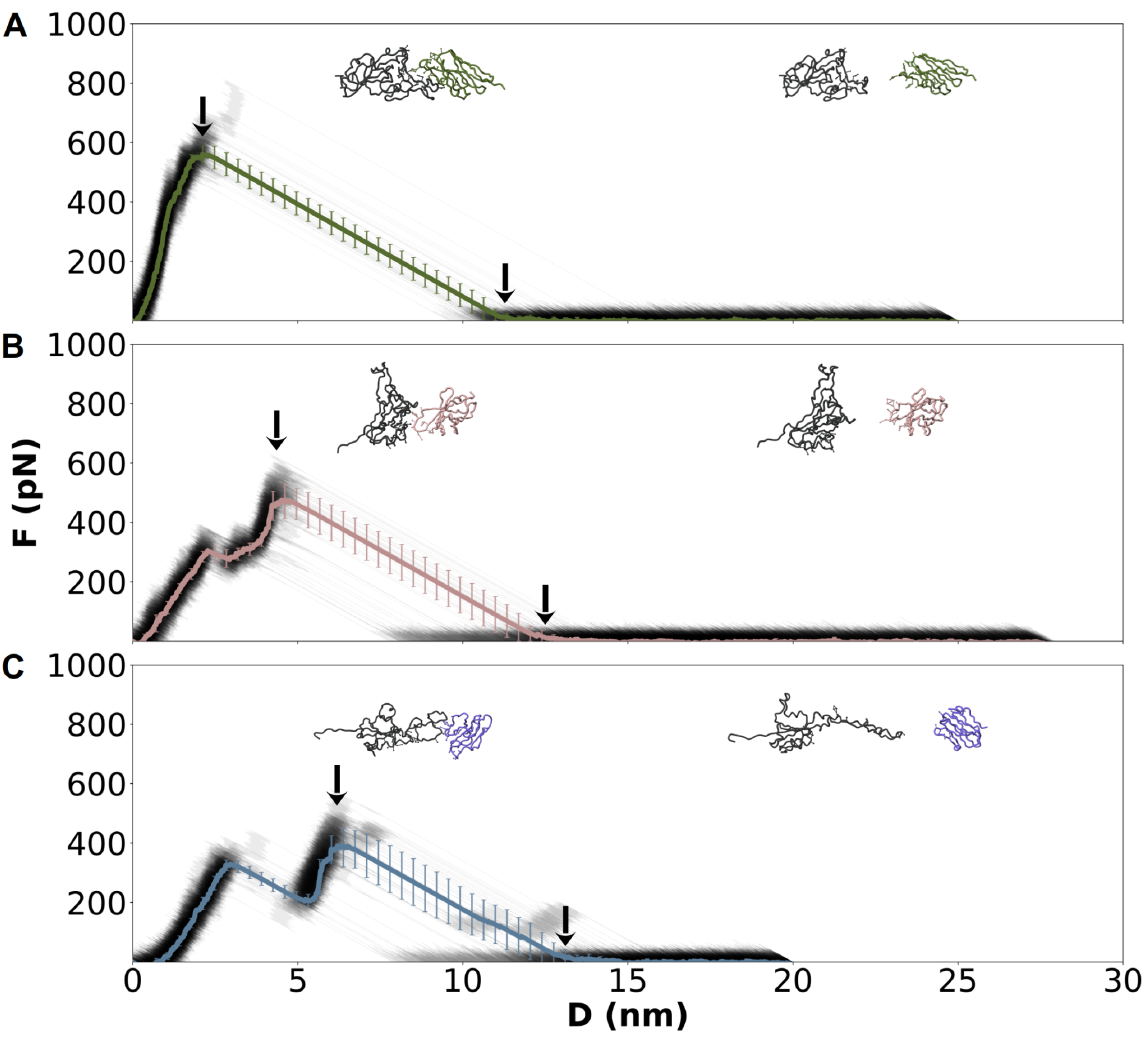
Mechanical response of Nb/RBD WT complexes under constant-velocity pulling. Force–displacement profiles are shown for (A) R14, (B) C1, and (C) n3113.1 (gray traces, mean in color). Insets display representative structures at the maximum force (F_max_, left arrow) and after dissociation (right arrow). Error bars denote standard deviations across 50 independent trajectories.

Across all systems, the L chain alone showed a remarkably consistent mechanical response that is independent of epitope geometry. This suggests that the L chain provides a baseline stabilizing role but cannot withstand large mechanical forces. In contrast, the H chain acts as the principal conduit for force transmission, channeling and maximizing the applied load through the RBD interface. Although less efficient on its own, its main role is amplified by the L chain, which reinforces and stabilizes the overall complex. Together, these findings highlight that Ab mechanical performance is not simply the sum of individual chains. Instead, the natural two-chain architecture evolved to integrate asymmetric yet complementary roles, with the H chain transmitting mechanical force and the L chain reinforcing stability through inter-chain cooperation. This arrangement was likely shaped by evolutionary pressures favoring both versatility and robustness ^59^.

### 3.4 Nanomechanics of Nb/RBD complexes

The binding affinity and neutralization potency of Nbs have been extensively characterized, with several studies reporting high stability and strong neutralization across SARS-CoV-2 variants ^60,61^. However, their mechanical behavior during force-induced dissociation remains largely unexplored to date ^12^. In Nbs, the absence of an L chain simplifies the architecture but also raises the question of whether their compact design transmits force more directly to the RBD interface, potentially altering the dissociation pathway compared with Abs.

#### 3.4.1 R14/RBD complex

The R14 Nb also targets the RBM and engages in interactions through CDRs 1–3 ^32^. Similar to PDI-231, it can only bind to the RBD in the up conformation. R14 demonstrates ultrahigh affinity, with no detectable dissociation observed for the WT, Alpha, Kappa, and Delta variants, indicating a very stable interaction. However, its neutralizing potency is reduced against Omicron variants, with measured K_D_ values ranging from 0.57 nM to 43 nM, reflecting the impact of RBM mutations on binding affinity ^32^.

We examined the mechanical dissociation of the RBD/R14 complex (PDB ID: 7WD1) for the WT, BA.4, and JN.1 variants. In all three cases, the dissociation proceeded via a rigid-body mechanism, with no significant unfolding of either the Nb or the RBD during mechanical dissociation. This consistent behavior suggests a mechanically stable binding mode across variants (Figure 5A and Table S5).

Despite the similar dissociation pathway, we observed pronounced differences in the maximum rupture forces (F_max_) among the variants (Table S11). The WT complex exhibited an average F_max_ of 596 ± 56 pN, indicative of a large mechanical stability at the R14/RBD interface. The BA.4 variant showed an even higher F_max_ in the range of 648.6 ± 44 pN (Figure S9A), pointing to enhanced mechanical stability despite its mutations. This increased resistance may reflect subtle changes in contact geometry or reinforcement of the binding interface introduced by BA.4-specific mutations (Tables S1 and S5).

In contrast, the JN.1 variant exhibited a markedly reduced F_max_ of 346 ± 35 pN (Figure 6A). This pronounced loss of mechanical stability implies that mutations in JN.1 compromise key stabilizing interactions at the R14/RBD interface, leading to premature rupture under mechanical forces. Importantly, the observation of a consistent rigid-body dissociation mode across all variants indicates that these mutations primarily modulate the strength of interfacial contacts, rather than altering the fundamental detachment mechanism.

**Figure 6.**
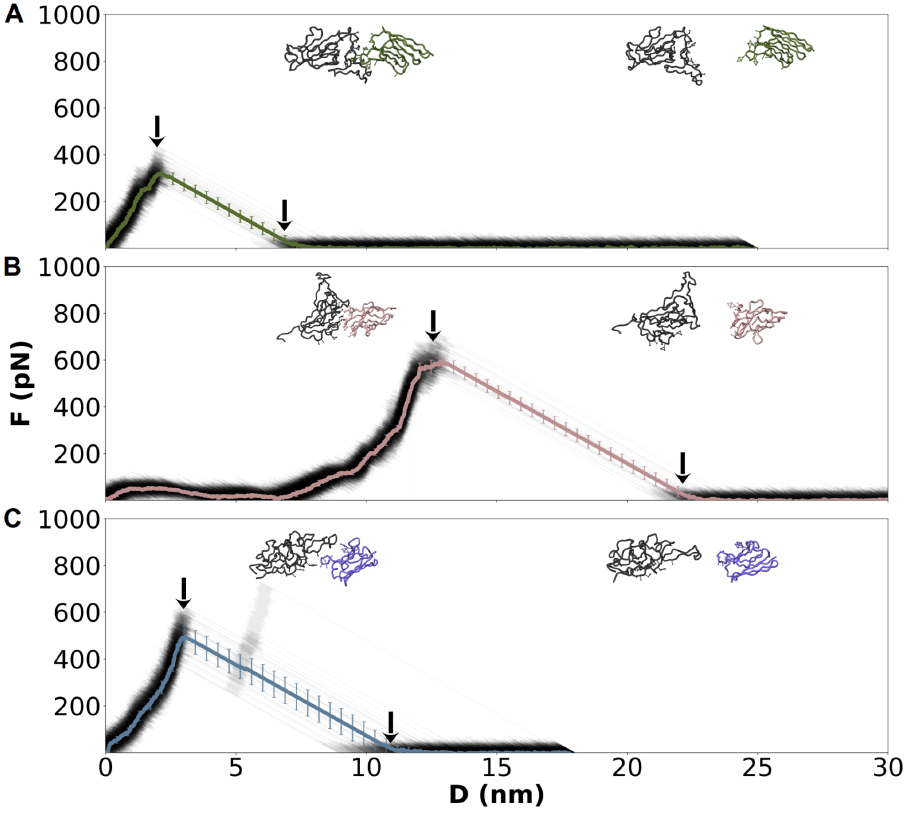
Mechanical response of Nb/RBD JN.1 complexes under constant-velocity pulling. Same as Figure 5.

#### 3.4.2 C1/RBD complex

Nb C1 exhibits high-affinity binding to the RBD of WT, Alpha, and Beta variants, with K_D_ in the 615–725 pM range ^33^. This Nb does not occlude the hACE2 binding site. Instead, it sterically hinders hACE2 binding, neutralizing the virus by occupying spatially adjacent regions (see Figure 1). C1 targets an epitope partially overlapping with that of S2X259, and its binding is associated with destabilization of the spike protein ^33^. This destabilization likely contributes to neutralization by disrupting the structural arrangement of the RBD that normally facilitates ACE2 binding, thereby impairing receptor engagement ^33^.

For C1 in the WT system, all trajectories exhibited a single prominent dissociation peak with an F_max_ of 479 ± 89, preceded by a characteristic drop in the mechanical force before reaching F_max_ (Figure 5B). Structurally, the RBD unfolded between residues A520 and S530, clearly distant from the RBM region. The process followed a rigid-body dissociation mechanism. Notably, the CDR3 loop was consistently the last element to disengage from the RBD, consistent with its deep insertion and tight interaction in the epitope pocket (Table S6).

In the BA.4 system, the mechanical response showed a large heterogeneity. Most CG-SMD trajectories exhibited a single dissociation peak with an F_max_ of 513±62 pN, typically preceded by a minor drop in force, similar to the WT pattern (Figure 5B and S9B). These cases involved partial unfolding of RBD residues F515–K528, followed by rigid-body dissociation of the Nb–antigen complex. However, in approx. 30% of the trajectories, a single dissociation peak occurred without unfolding of either the RBD or the Nb. This outcome indicates that, in certain configurations, interfacial contacts can be disrupted directly, without any preceding structural rearrangement, highlighting an alternative dissociation pathway unique to BA.4.

For the JN.1 variant, the mechanical profiles also showed a single dissociation peak with an F_max_ of 603 ± 56. Most curves build up slowly and converge at a single, sharply defined rupture event (Figure 6B). This suggests a more flexible binding interface in the JN.1 system, potentially due to structural rearrangements in the RBD or reduced initial contact strength. Despite this, dissociation remained rigid-body in nature, and unfolding of residues A520–S530 in the RBD consistently preceded the rupture. Notably, no unfolding was detected in the Nb, underscoring its mechanical stability throughout the process.

#### 3.4.3 n3113.1/RBD complex

We analyzed the mechanical dissociation of the RBD–n3113.1 complex for the WT, BA.4, and JN.1 variants. For the WT system, the dissociation profiles followed more complex pathways predominantly characterized by a sawtooth pattern, related to the sequential unfolding of the RBD (Figure 5C). In most SMD trajectories, we observed unfolding of the RBD in two regions: partial unfolding of its C-terminal segment (residues L517–K528) followed by more substantial structural loss in the N437–P507 region (Table S7). Despite this partial unfolding, the Nb itself remained folded and dissociated as a rigid-body. In a subset of seven trajectories, we observed a simpler dissociation pattern featuring a single force peak, accompanied by only minor unfolding of the RBD C-terminus (L517–K528), again with rigid-body detachment of n3113.1.

The BA.4 variant also predominantly exhibited a single-peak dissociation profile, indicative of rigid-body detachment. In the majority of SMD trajectories, both the RBD and the Nb remained fully folded throughout the pulling process (Figure S9C). In some cases, the presence of a force bump precedes the main rupture peak, associated with subtle unfolding of RBD residues L517–K528. Nevertheless, these events did not alter the global rigid-body nature of the Nb detachment. In contrast, the JN.1 variant displayed a simpler mechanical response (Figure 6C). Most dissociation events featured a single peak consistent with rigid-body rupture, without significant unfolding of either the RBD or the Nb. In a few trajectories, slight unfolding of the RBD C-terminal region (residues L517–K528) was observed before dissociation, but the Nb consistently dissociated without structural deformation.

## 4 Conclusions

Our structure-based methodology extends the Martini 3 force field ^26^ to study large protein conformational changes, capturing both the stability of protein complexes at equilibrium ^12,21^ and the subtle differences between RBD variants in complex with Abs and Nbs, particularly the partial unfolding of the RBD under mechanical forces. To improve the accuracy of GōMartini 3, this study optimized the contacts using AA-MD simulations. GōMartini 3 simulations require a well-defined reference structure to define the parameters underlying secondary structure. In contrast, other protein models, such as the OLIVES approach ^62^ in Martini 3, or pragmatic CG models like SIRAH and UNRES ^20^ do not rely on a reference structure for secondary structure parametrization, but they have not been tested under mechanical forces. The scalability of GōMartini 3 simulations for oligomeric systems was initially limited by the definition of Gō contacts between chains, but this issue was recently addressed by Korshunova et al. for Martini 3 ^63^.

Our analysis highlights fundamental differences in the dissociation behavior of Abs and Nbs bound to the SARS-CoV-2 RBD under mechanical force, revealing how structural architecture, epitope location, and viral mutations influence mechanical resilience. Across all complexes studied, dissociation of processes occurred predominantly through rigid-body mechanisms. However, unfolding events, both in the RBD and in Ab chains, emerged in a variant- and chain-pulling–dependent manner. In Ab/RBD complexes, pulling from different chains frequently led to asymmetric responses. For instance, pulling from the H chain of PDI-231 or S2X259 often induces limited unfolding in the Ab’s constant region or in RBD residues 521–529. In contrast, pulling from the L chain promoted sequential unbinding, sometimes accompanied by unfolding of the L chain’s C-terminal region. Notably, the JN.1 variant tended to display reduced unfolding and more simultaneous chain detachment, suggesting enhanced inter-chain coordination or altered mechanical rigidity. A comparison of single-chain and intact Ab systems further demonstrated that the H chain provides the dominant load-bearing capacity (~230–430 pN), while the L chain sustains only ~200 pN across epitopes. Importantly, their combination yields higher rupture forces than either chain alone, reflecting cooperative stabilization and distributed force transmission unique to the dual-chain architecture.

Nb complexes, by contrast, exhibited greater mechanical uniformity. R14 and C1 remained structurally rigid during dissociation, with unfolding restricted largely to RBD residues 515–529. C1 dissociation was often preceded by a small force peak before F_max_, consistent with partial C-terminal RBD unfolding, whereas R14, despite targeting a similar epitope to PDI-231, consistently achieved higher rupture forces, particularly in BA.4. This indicates that the compact, single-domain geometry of Nbs facilitates more efficient force transmission and better resistance to mechanical stress compared to Abs. In JN.1.

In non-hACE2-competing complexes (R1-32/RBD and n3113.1/RBD), mechanical dissociation was dominated by unfolding of the RBD within the labile S438–P507 region, whereas the Ab and Nb scaffolds retained their structural integrity. This region overlaps with a β-sheet–loop–β-sheet network identified in prior pulling studies as a primary rupture hotspot^48^, stabilized by contacts involving A348–Y351 and V401, L452, Y451, Y453, R454, and L492. Disruption of these interactions initiates β-strand separation and destabilization of the RBD core, explaining the recurrent unfolding observed in our simulations. Variant-specific effects were evident, with BA.4 frequently enhancing mechanical resistance and JN.1 reducing it across multiple complexes. Importantly, the structural plasticity of the RBD in regions 516–529 and 438–507 emerges as a common mechanical fracture point for both Abs and Nbs.

Our findings underscore the value of mechanical resilience as a complementary parameter to binding affinity in therapeutic design. Mapping and exploiting mechanically sensitive regions, particularly conserved contact clusters within the RBD, may guide the development of next-generation Abs and Nbs with improved stability against structural destabilization and immune escape by emerging variants.

## Author contributions

L. F. C. V.: writing – review & editing, writing – original draft, methodology, investigation, formal analysis. G. E. O. R.: review, scientific discussion,. A. B. P.: writing – review & editing, writing – original draft, supervision, methodology, investigation, funding acquisition, conceptualization. S.-J. M.: writing – review & editing, scientific discussion..

## Data availability

The data supporting the findings of this study are openly available in the Zenodo repository: https://zenodo.org/records/15608522. The repository includes all input files and simulations utilized in this research:

## Acknowledgements

A.B.P. acknowledges financial support from the National Science Center, Poland, under grant 2022/45/B/NZ1/02519 and gratefully acknowledges Polish high-performance computing infrastructure PLGrid (HPC Center: ACK Cyfronet AGH) for providing computer facilities and support within computational grant no. PLG/2024/017332 and PLG/2025/018510. This work was financed in part by DGAPA, UNAM [PAPIIT, IN209624].

## References

1 W. Msemburi, A. Karlinsky, V. Knutson, S. Aleshin-Guendel, S. Chatterji and J. Wakefield, Nature, 2023, 613, 130–137.

2 S. Zhou, P. Lv, M. Li, Z. Chen, H. Xin, S. Reilly and X. Zhang, Biomed. Pharmacother., 2023, 159, 114242.

3 A. Sternberg and C. Naujokat, Life Sci., 2020, 257, 118056.

4 J. Zhang, T. Xiao, Y. Cai and B. Chen, Curr. Opin. Virol., 2021, 50, 173–182.

5 J. Lan, J. Ge, J. Yu, S. Shan, H. Zhou, S. Fan, Q. Zhang, X. Shi, Q. Wang, L. Zhang and X. Wang, Nature, 2020, 581, 215–220.

6 A. Deshpande, B. D. Harris, L. Martinez-Sobrido, J. J. Kobie and M. R. Walter, Front. Immunol., 2021, 12, 691715.

7 A. Ray, T. T. Minh Tran, R. D. Santos Natividade, R. A. Moreira, J. D. Simpson, D. Mohammed, M. Koehler, S. J. L Petitjean, Q. Zhang, F. Bureau, L. Gillet, A. B. Poma and D. Alsteens, ACS Nanosci. Au, 2024, 4, 136–145.

8 G. E. Olivos-Ramirez, L. F. Cofas-Vargas, T. Madl and A. B. Poma, Pathogens,, DOI:10.3390/pathogens14030274.

9 S. Kim, Y. Liu, M. Ziarnik, Y. Cao, X. F. Zhang and W. Im, bioRxivorg, 2022.

10 D. Koirala, P. M. Yangyuoru and H. Mao, Rev. Anal. Chem.,, DOI:10.1515/revac-2013-0004.

11 W. Liu, D. Tang, X.-X. Xu, Y.-J. Liu and Y. Jiu, Front. Bioeng. Biotechnol., 2021, 9, 764516.

12 L. F. Cofas-Vargas, G. E. Olivos-Ramirez, M. Chwastyk, R. A. Moreira, J. L. Baker, S. J. Marrink and A. B. Poma, Nanoscale, 2024, 16, 18824–18834.

13 M. Golcuk, A. Hacisuleyman, S. Z. Yilmaz, E. Taka, A. Yildiz and M. Gur, J. Chem. Inf. Model., 2022, 62, 2490–2498.

14 M. Golcuk, A. Hacisuleyman, B. Erman, A. Yildiz and M. Gur, J. Chem. Inf. Model., 2021, 61, 5152–5160.

15 H. Nguyen and M. S. Li, Sci. Rep., 2022, 12, 1–15.

16 S. Muyldermans and V. V. Smider, Curr. Opin. Immunol., 2016, 40, 7–13.

17 B.-K. Jin, S. Odongo, M. Radwanska and S. Magez, Int. J. Mol. Sci., 2023, 24, 5994.

18 J. Huo, A. Le Bas, R. R. Ruza, H. M. E. Duyvesteyn, H. Mikolajek, T. Malinauskas, T. K. Tan, P. Rijal, M. Dumoux, P. N. Ward, J. Ren, D. Zhou, P. J. Harrison, M. Weckener, D. K. Clare, V. K. Vogirala, J. Radecke, L. Moynié, Y. Zhao, J. Gilbert-Jaramillo, M. L. Knight, J. A. Tree, K. R. Buttigieg, N. Coombes, M. J. Elmore, M. W. Carroll, L. Carrique, P. N. M. Shah, W. James, A. R. Townsend, D. I. Stuart, R. J. Owens and J. H. Naismith, Nat. Struct. Mol. Biol., 2020, 27, 846–854.

19 D. K. Agarwal, V. Nandwana, S. E. Henrich, V. P. V. N. Josyula, C. S. Thaxton, C. Qi, L. M. Simons, J. F. Hultquist, E. A. Ozer, G. S. Shekhawat and V. P. Dravid, Biosens. Bioelectron., 2022, 195, 113647.

20 A. B. Poma, A. Hinostroza Caldas, L. F. Cofas-Vargas, M. S. Jones, A. L. Ferguson and L. Medrano Sandonas, Biophys J,, DOI:10.1016/j.bpj.2025.06.019.

21 Z. Liu, R. A. Moreira, A. Dujmović, H. Liu, B. Yang, A. B. Poma and M. A. Nash, Nano Lett., 2022, 22, 179–187.

22 J. Alegre-Cebollada, Biophys. Rev., 2021, 13, 435–454.

23 A. B. Poma, T. T. M. Thu, L. T. M. Tri, H. L. Nguyen and M. S. Li, J. Phys. Chem. B, 2021, 125, 7628–7637.

24 M. C. R. Melo, D. E. B. Gomes and R. C. Bernardi, J. Am. Chem. Soc., 2023, 145, 70–77.

25 P. C. T. Souza, L. Borges-Araújo, C. Brasnett, R. A. Moreira, F. Grünewald, P. Park, L. Wang, H. Razmazma, A.C. Borges-Araújo, L. F. Cofas-Vargas, L. Monticelli, R. Mera-Adasme, M. N. Melo, S. Wu, S. J. Marrink, A. B. Poma and S. Thallmair, Nat Commun, 2025, 16, 4051.

26 P. C. T. Souza, R. Alessandri, J. Barnoud, S. Thallmair, I. Faustino, F. Grünewald, I. Patmanidis, H. Abdizadeh, B. M. H. Bruininks, T. A. Wassenaar, P. C. Kroon, J. Melcr, V. Nieto, V. Corradi, H. M. Khan, J. Domański, M. Javanainen, H. Martinez-Seara, N. Reuter, R. B. Best, I. Vattulainen, L. Monticelli, X. Periole, D. P. Tieleman, A. H. de Vries and S. J. Marrink, Nat Methods, 2021, 18, 382–388.

27 A. B. Poma, M. Cieplak and P. E. Theodorakis, J. Chem. Theory Comput., 2017, 13, 1366–1374.

28 K. Wołek, À. Gómez-Sicilia and M. Cieplak, J. Chem. Phys., 2015, 143, 243105.

29 A. K. Wheatley, P. Pymm, R. Esterbauer, M. H. Dietrich, W. S. Lee, D. Drew, H. G. Kelly, L.-J. Chan, F. L. Mordant, K. A. Black, A. Adair, H.-X. Tan, J. A. Juno, K. M. Wragg, T. Amarasena, E. Lopez, K. J. Selva, E. R. Haycroft, J. P. Cooney, H. Venugopal, L. L. Tan, M. T. O Neill, C. C. Allison, D. Cromer, M. P. Davenport, R. A. Bowen, A. W. Chung, M. Pellegrini, M. T. Liddament, A. Glukhova, K. Subbarao, S. J. Kent and W.-H. Tham, Cell Rep., 2021, 37, 109822.

30 T. N. Starr, N. Czudnochowski, Z. Liu, F. Zatta, Y.-J. Park, A. Addetia, D. Pinto, M. Beltramello, P. Hernandez, A. J. Greaney, R. Marzi, W. G. Glass, I. Zhang, A. S. Dingens, J. E. Bowen, M. A. Tortorici, A. C. Walls, J. A. Wojcechowskyj, A. De Marco, L. E. Rosen, J. Zhou, M. Montiel-Ruiz, H. Kaiser, J. R. Dillen, H. Tucker, J. Bassi, C. Silacci-Fregni, M. P. Housley, J. di Iulio, G. Lombardo, M. Agostini, N. Sprugasci, K. Culap, S. Jaconi, M. Meury, E. Dellota Jr, R. Abdelnabi, S.-Y. C. Foo, E. Cameroni, S. Stumpf, T. I. Croll, J. C. Nix, C. Havenar-Daughton, L. Piccoli, F. Benigni, J. Neyts, A. Telenti, F. A. Lempp, M. S. Pizzuto, J. D. Chodera, C. M. Hebner, H. W. Virgin, S. P. J. Whelan, D. Veesler, D. Corti, J. D. Bloom and G. Snell, Nature, 2021, 597, 97–102.

31 P. He, B. Liu, X. Gao, Q. Yan, R. Pei, J. Sun, Q. Chen, R. Hou, Z. Li, Y. Zhang, J. Zhao, H. Sun, B. Feng, Q. Wang, H. Yi, P. Hu, P. Li, Y. Zhang, Z. Chen, X. Niu, X. Zhong, L. Jin, X. Liu, K. Qu, K. A. Ciazynska, A. P. Carter, J. A. G. Briggs, J. Chen, J. Liu, X. Chen, J. He, L. Chen and X. Xiong, Nat. Microbiol., 2022, 7, 1635–1649.

32 H. Liu, L. Wu, B. Liu, K. Xu, W. Lei, J. Deng, X. Rong, P. Du, L. Wang, D. Wang, X. Zhang, C. Su, Y. Bi, H. Chen, W. J. Liu, J. Qi, Q. Cui, S. Qi, R. Fan, J. Jiang, G. Wu, G. F. Gao and Q. Wang, Cell Rep. Med., 2023, 4, 100918.

33 J. Huo, H. Mikolajek, A. Le Bas, J. J. Clark, P. Sharma, A. Kipar, J. Dormon, C. Norman, M. Weckener, D. K. Clare, P. J. Harrison, J. A. Tree, K. R. Buttigieg, F. J. Salguero, R. Watson, D. Knott, O. Carnell, D. Ngabo, M. J. Elmore, S. Fotheringham, A. Harding, L. Moynié, P. N. Ward, M. Dumoux, T. Prince, Y. Hall, J. A. Hiscox, A. Owen, W. James, M. W. Carroll, J. P. Stewart, J. H. Naismith and R. J. Owens, Nat. Commun., 2021, 12, 5469.

34 Z. Yang, Y. Wang, Y. Jin, Y. Zhu, Y. Wu, C. Li, Y. Kong, W. Song, X. Tian, W. Zhan, A. Huang, S. Zhou, S. Xia, X. Tian, C. Peng, C. Chen, Y. Shi, G. Hu, S. Du, Y. Wang, Y. Xie, S. Jiang, L. Lu, L. Sun, Y. Song and T. Ying, Signal Transduct. Target. Ther., 2021, 6, 378.

35 A. Sali and T. L. Blundell, J. Mol. Biol., 1993, 234, 779–815.

36 C. Tian, K. Kasavajhala, K. A. A. Belfon, L. Raguette, H. Huang, A. N. Migues, J. Bickel, Y. Wang, J. Pincay, Q. Wu and C. Simmerling, J Chem Theory Comput, 2020, 16, 528–552.

37 D. A. Case, H. M. Aktulga, K. Belfon, I. Y. Ben-Shalom, J. T. Berryman, S. R. Brozell, D. S. Cerutti, T. E. Cheatham, III, G. A. Cisneros, V. W. D. Cruzeiro, T. A. Darden, N. Forouzesh, M. Ghazimirsaeed, G. Giambaşu, T. Giese, M. K. Gilson, H. Gohlke, A. W. Goetz, J. Harris, Z. Huang, S. Izadi, S. A. Izmailov, K. Kasavajhala, M. C. Kaymak, A. Kovalenko, T. Kurtzman, T. S. Lee, P. Li, Z. Li, C. Lin, J. Liu, T. Luchko, R. Luo, M. Machado, M. Manathunga, K. M. Merz, Y. Miao, O. Mikhailovskii, G. Monard, H. Nguyen, K. A. O’Hearn, A. Onufriev, F. Pan, S. Pantano, A. Rahnamoun, D. R. Roe, A. Roitberg, C. Sagui, S. Schott-Verdugo, A. Shajan, J. Shen, C. L. Simmerling, N. R. Skrynnikov, J. Smith, J. Swails, R. C. Walker, J. Wang, J. Wang, X. Wu, Y. Wu, Y. Xiong, Y. Xue, D. M. York, C. Zhao, Q. Zhu, A. P. Kollman (2024), Amber, University of California and S. Francisco., Amber24, 2024.

38 R. Salomon-Ferrer, A. W. Götz, D. Poole, S. Le Grand and R. C. Walker, J Chem Theory Comput, 2013, 9, 3878–3888.

39 P. Eastman, J. Swails, J. D. Chodera, R. T. McGibbon, Y. Zhao, K. A. Beauchamp, L.-P. Wang, A. C. Simmonett, M. P. Harrigan, C. D. Stern, R. P. Wiewiora, B. R. Brooks and V. S. Pande, PLoS Comput Biol, 2017, 13, e1005659.

40 S. Izadi, R. Anandakrishnan and A. V. Onufriev, J. Phys. Chem. Lett., 2014, 5, 3863–3871.

41 A. C. Simmonett and B. R. Brooks, J. Chem. Phys., 2021, 154, 054112.

42 D. S. Cerutti, R. E. Duke, T. A. Darden and T. P. Lybrand, J. Chem. Theory Comput., 2009, 5, 2322.

43 D. J. Sindhikara, S. Kim, A. F. Voter and A. E. Roitberg, J. Chem. Theory Comput., 2009, 5, 1624–1631.

44 J. Åqvist, P. Wennerström, M. Nervall, S. Bjelic and B. O. Brandsdal, Chem. Phys. Lett., 2004, 384, 288–294.

45 J.-P. Ryckaert, G. Ciccotti and H. J. C. Berendsen, J. Comput. Phys., 1977, 23, 327–341.

46 M. R. Shirts, C. Klein, J. M. Swails, J. Yin, M. K. Gilson, D. L. Mobley, D. A. Case and E. D. Zhong, J. Comput. Aided Mol. Des., 2017, 31, 147–161.

47 C. W. Hopkins, S. Le Grand, R. C. Walker and A. E. Roitberg, J. Chem. Theory Comput., 2015, 11, 1864–1874.

48 R. A. Moreira, M. Chwastyk, J. L. Baker, H. V. Guzman and A. B. Poma, Nanoscale, 2020, 12, 16409–16413.

49 P. C. Kroon, F. Grunewald, J. Barnoud, M. van Tilburg, P. C. T. Souza, T. A. Wassenaar and S. J. Marrink, eLife, 2024.

50 R. P. Joosten, T. A. H. te Beek, E. Krieger, M. L. Hekkelman, R. W. W. Hooft, R. Schneider, C. Sander and G. Vriend, Nucleic Acids Res., 2011, 39, D411–9.

51 L. F. Cofas-Vargas, R. A. Moreira, S. Poblete, M. Chwastyk and A. B. Poma, Acta Phys. Pol. A, 2024, 145, S9–S9.

52 G. Bussi, D. Donadio and M. Parrinello, J. Chem. Phys., 2007, 126, 014101.

53 M. Bernetti and G. Bussi, J. Chem. Phys., 2020, 153, 114107.

54 M. J. Abraham, T. Murtola, R. Schulz, S. Páll, J. C. Smith, B. Hess and E. Lindahl, SoftwareX, 2015, 1-2, 19–25.

55 M. A. Tortorici, N. Czudnochowski, T. N. Starr, R. Marzi, A. C. Walls, F. Zatta, J. E. Bowen, S. Jaconi, J. Di Iulio, Z. Wang, A. De Marco, S. K. Zepeda, D. Pinto, Z. Liu, M. Beltramello, I. Bartha, M. P. Housley, F. A. Lempp, L. E. Rosen, E. Dellota Jr, H. Kaiser, M. Montiel-Ruiz, J. Zhou, A. Addetia, B. Guarino, K. Culap, N. Sprugasci, C. Saliba, E. Vetti, I. Giacchetto-Sasselli, C. S. Fregni, R. Abdelnabi, S.-Y. C. Foo, C. Havenar-Daughton, M. A. Schmid, F. Benigni, E. Cameroni, J. Neyts, A. Telenti, H. W. Virgin, S. P. J. Whelan, G. Snell, J. D. Bloom, D. Corti, D. Veesler and M. S. Pizzuto, Nature, 2021, 597, 103–108.

56 Q. He, L. Wu, Z. Xu, X. Wang, Y. Xie, Y. Chai, A. Zheng, J. Zhou, S. Qiao, M. Huang, G. Shang, X. Zhao, Y. Feng, J. Qi, G. F. Gao and Q. Wang, Cell Rep. Med., 2023, 4, 100991.

57 H. Ma, C. Ó’Fágáin and R. O’Kennedy, Biochimie, 2020, 177, 213–225.

58 M. Li, F. Lou and H. Fan, Signal Transduct. Target. Ther., 2022, 7, 28.

59 P. Dudzic, D. Chomicz, W. Bielska, I. Jaszczyszyn, M. Zieliński, B. Janusz, S. Wróbel, M.-M. L. Pannérer, A. Philips, P. Ponraj, S. Kumar and K. Krawczyk, Commun. Biol., 2025, 8, 1110.

60 D. Wrapp, D. De Vlieger, K. S. Corbett, G. M. Torres, N. Wang, W. Van Breedam, K. Roose, L. van Schie, VIB-CMB COVID-19 Response Team, M. Hoffmann, S. Pöhlmann, B. S. Graham, N. Callewaert, B. Schepens, X. Saelens and J. S. McLellan, Cell, 2020, 181, 1004–1015.e15.

61 Y. Xiang, S. Nambulli, Z. Xiao, H. Liu, Z. Sang, W. P. Duprex, D. Schneidman-Duhovny, C. Zhang and Y. Shi, Science, 2020, 370, 1479–1484.

62 K. B. Pedersen, L. Borges-Araújo, A. D. Stange, P. C. T. Souza, S. J. Marrink and B. Schiøtt, J Chem Theory Comput,, DOI:10.1021/acs.jctc.4c00553.

63 K. Korshunova, J. Kiuru, J. Liekkinen, G. Enkavi, I. Vattulainen and B. M. H. Bruininks, J Chem Theory Comput, 2024, 20, 7635–7645.

