## Supplementary Information for "A Comparative Nanomechanical Study of Antibody and Nanobody Binding to SARS-CoV-2 Variants"

**This file includes:**

Supplementary Figures S1 to S19

Tables S1-S11

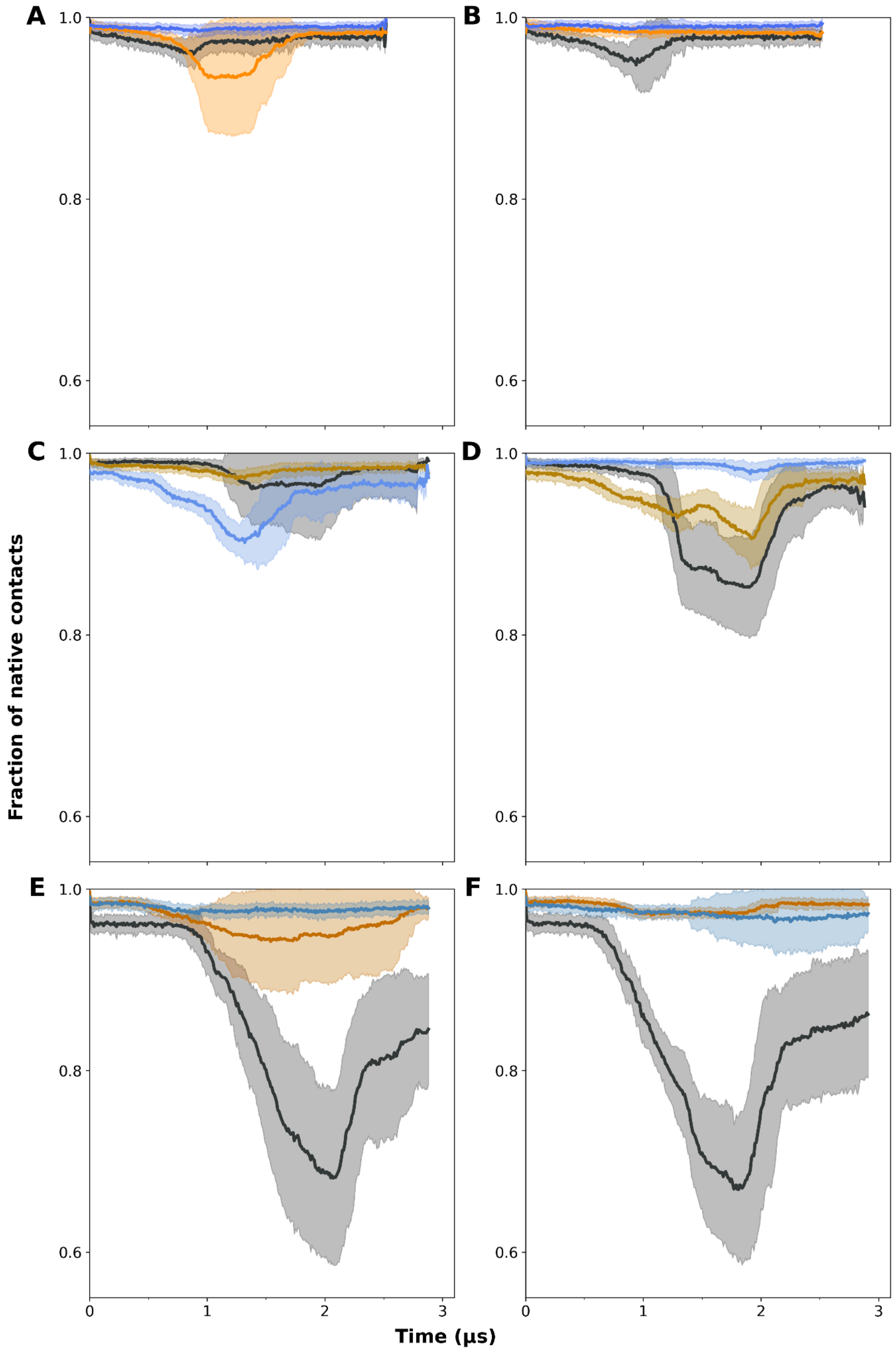


**Figure S1. Contact frequency profiles over simulation time for RBD/Ab WT complexes.** Timeline of the fraction of native contacts for the RBD and Ab H and L chains. RBD is shown in gray, H chains in orange/yellow, and L chains in blue. Panels in the left column (A, C, E) show SMD pulling from the H chain, and panels in the right column (B, D, F) show pulling from the L chain. Each row corresponds to a different complex: row 1, RBD/PDI-231 WT; row 2, S2XS259; row 3, R1-32.

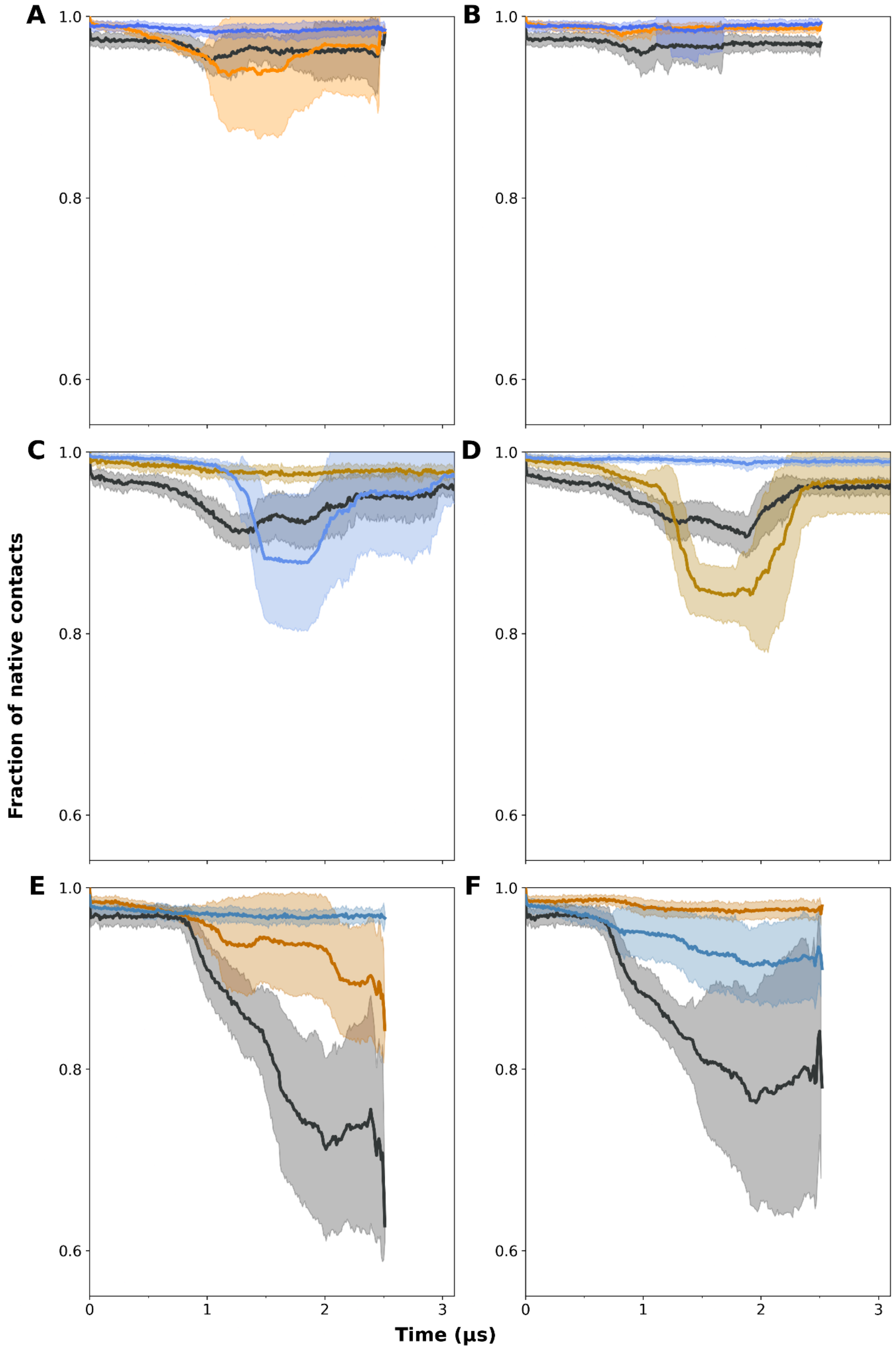


**Figure S2. Contact frequency profiles over simulation time for RBD/Ab BA.4 complexes.** Same as Figure S1.

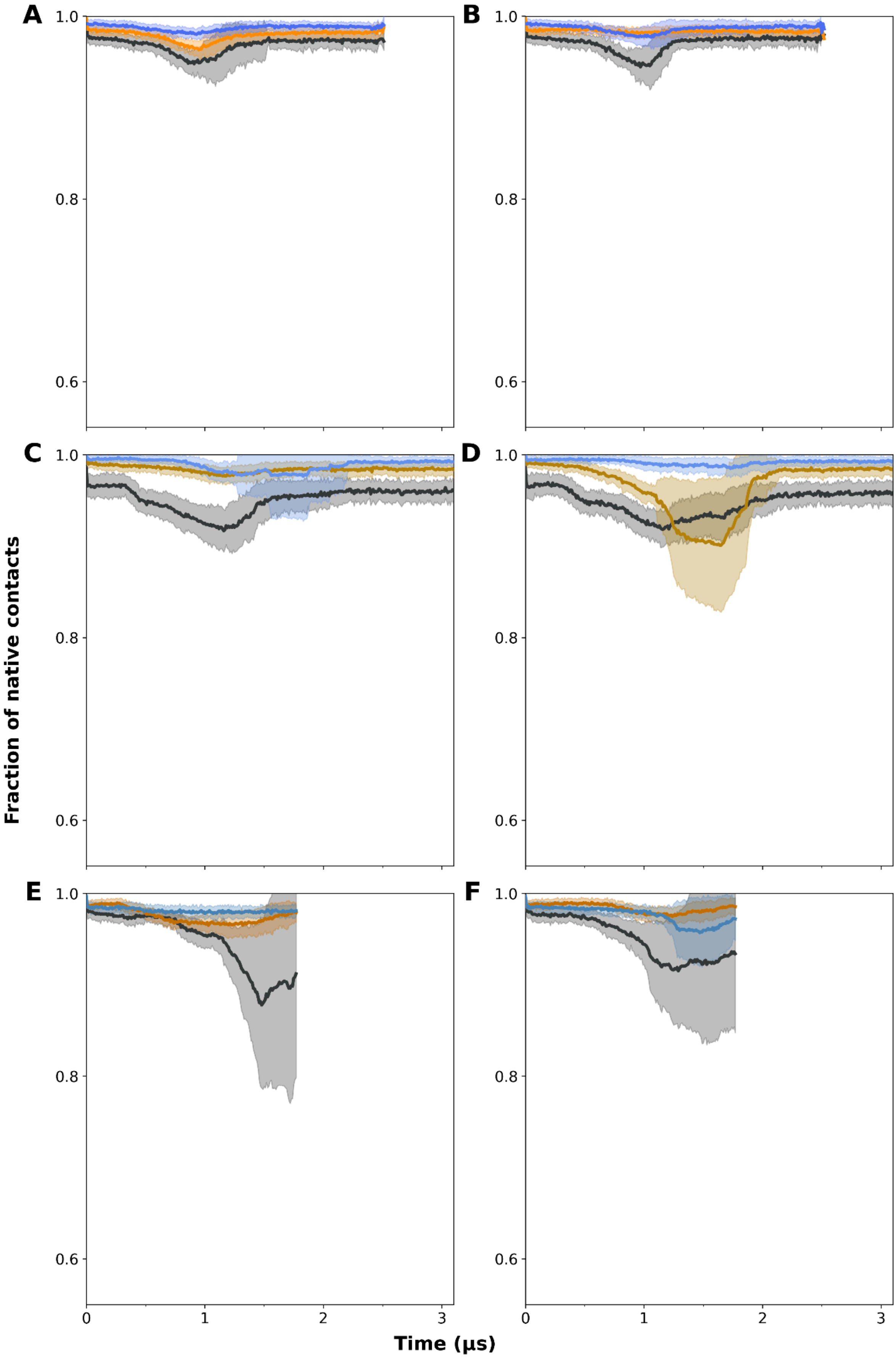


**Figure S3. Contact frequency profiles over simulation time for RBD/Ab JN.1 complexes.** Same as Figure S1

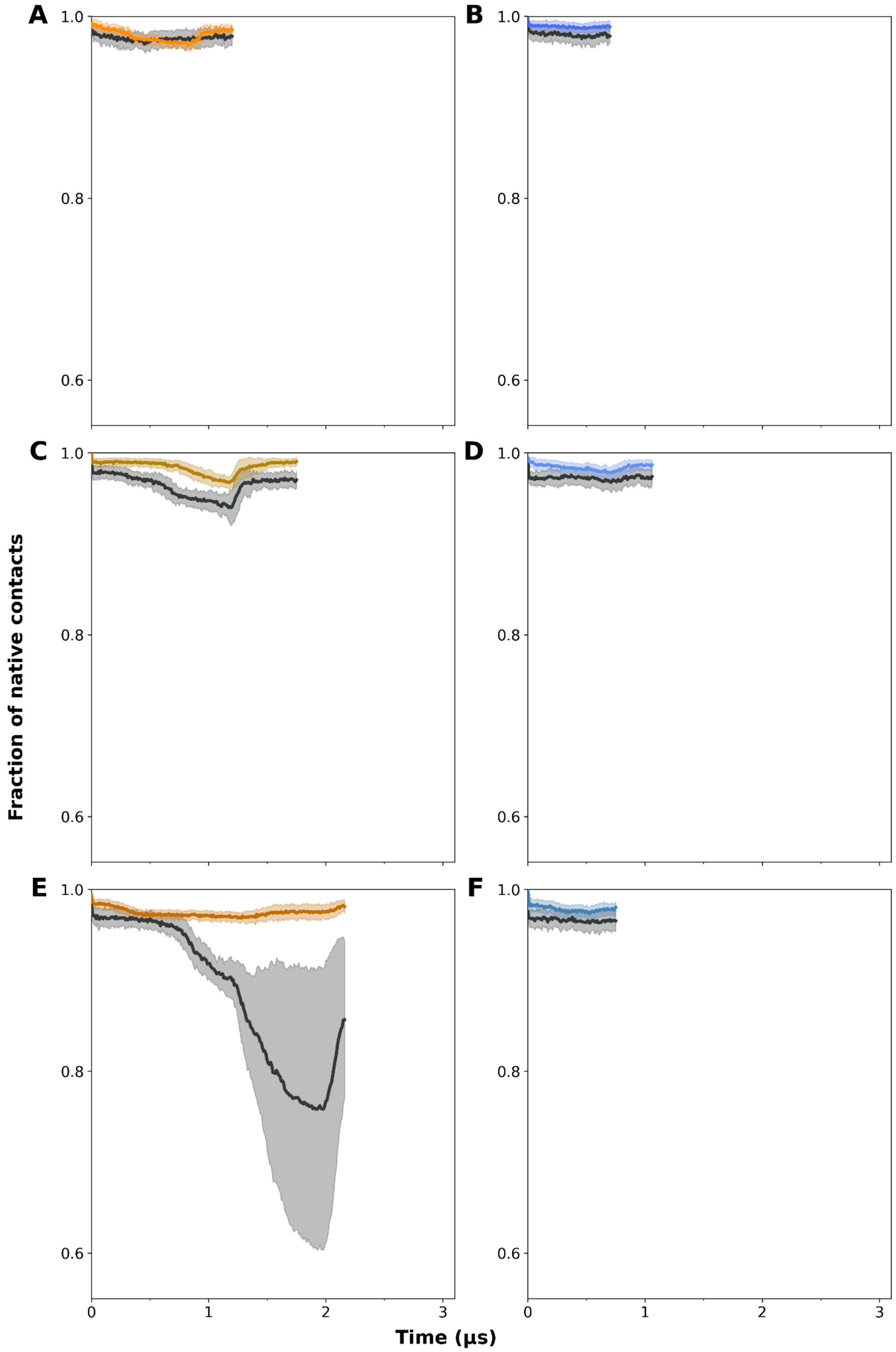


**Figure S4. Mechanical response of Contact frequency profiles over simulation time for the Ab/RBD WT complexes under constant-velocity pulling from individual chains.** Force-displacement profiles are shown of (A, B) PDI-231, (C, D) S2X259, and (E, F) R1-32. Pulling was applied from the H chain (left column) or the L chain (right column). Solid lines correspond to the mean response.

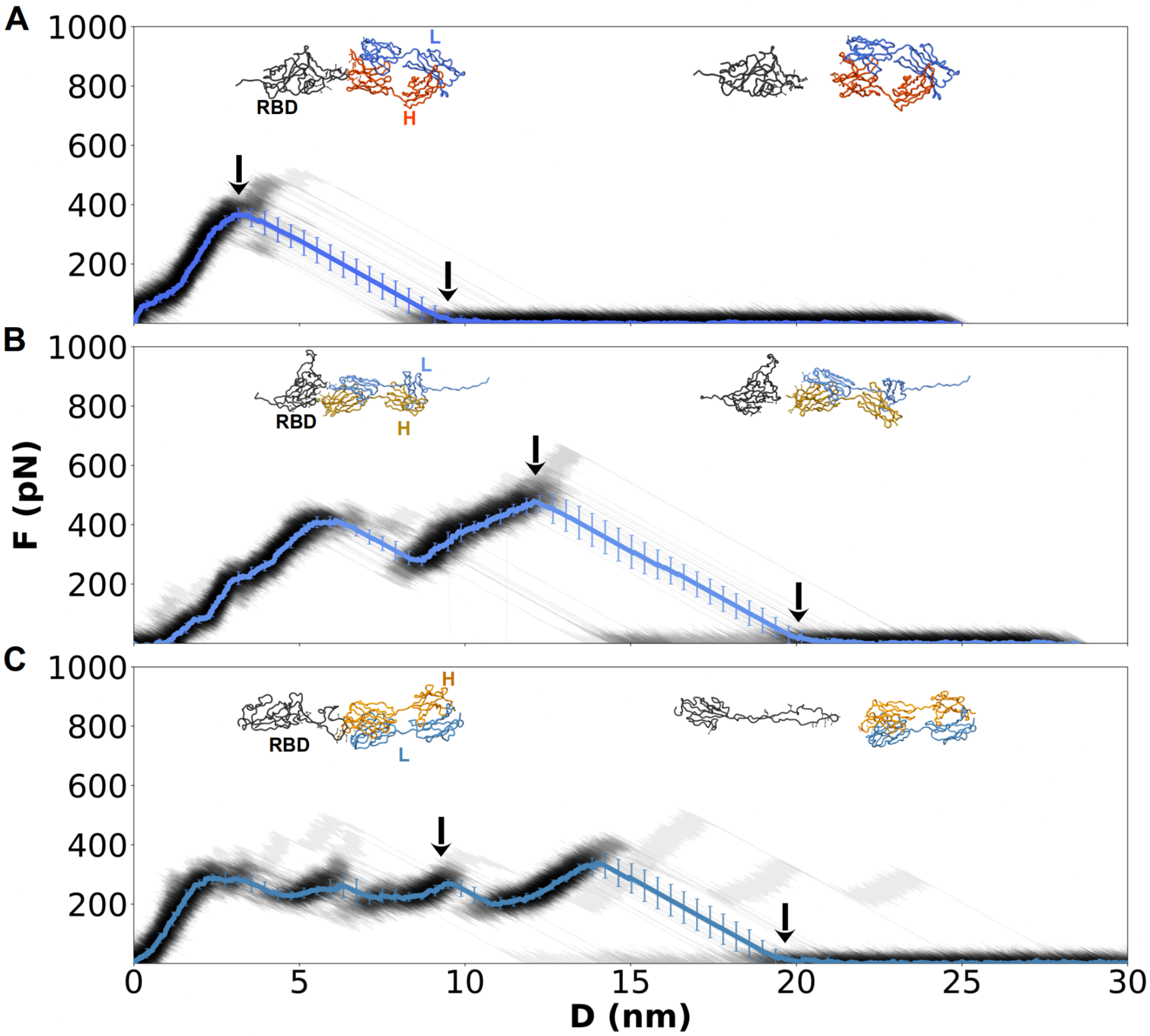


**Figure S5. Mechanical response of Ab/RBD WT complexes under constant-velocity pulling from the L chain.** Force–displacement profiles are shown for (A) PDI-231, (B) S2X259, and (C) R1-32 (gray traces, mean in color). Insets display representative structures at the maximum force (Fmax, left arrow) and after dissociation (right arrow). Error bars denote standard deviations across 50 independent trajectories.

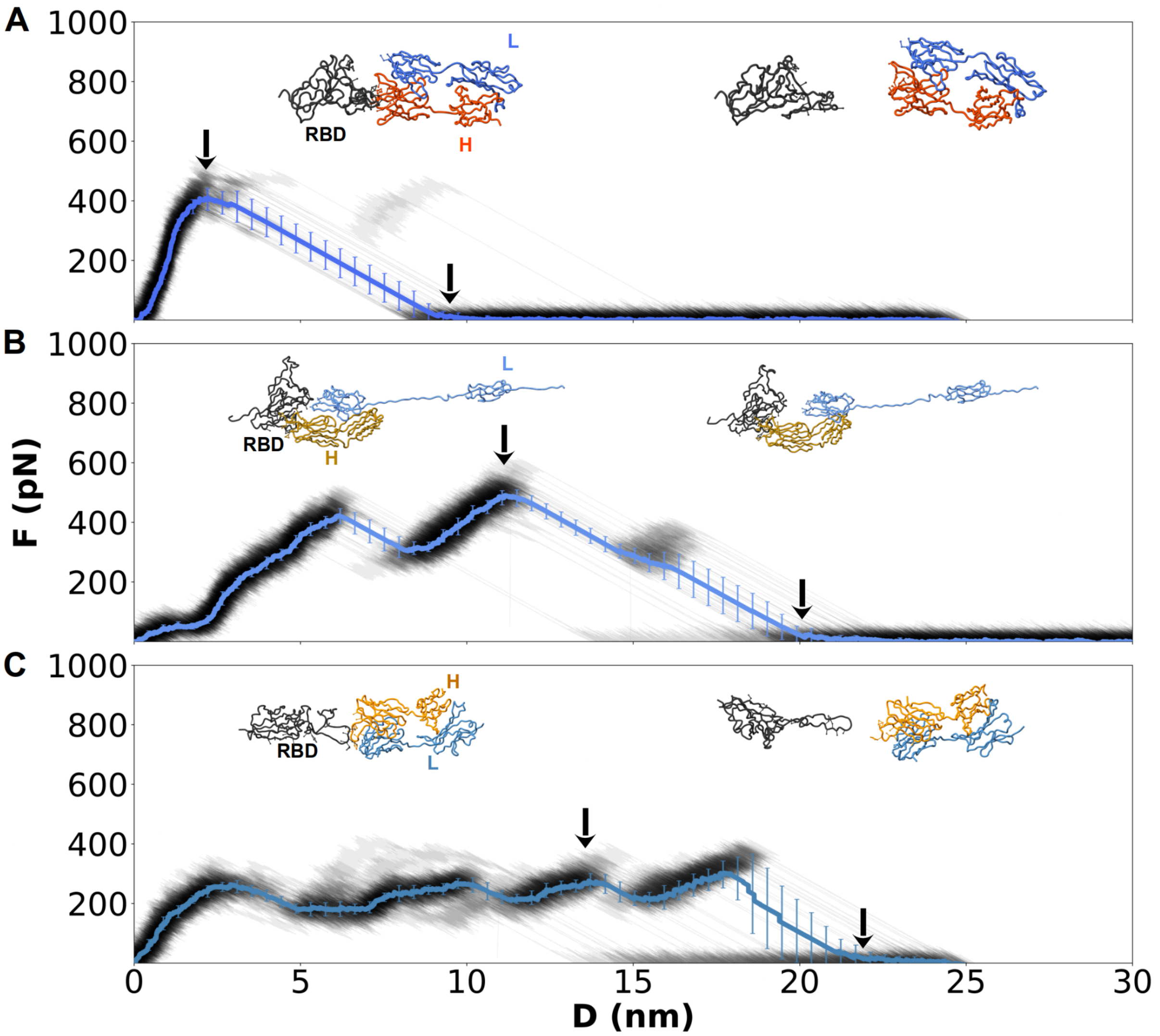


**Figure S6. Mechanical response of Ab/RBD BA.4 complexes under constant-velocity pulling from the L chain.** Same as Figure S4.

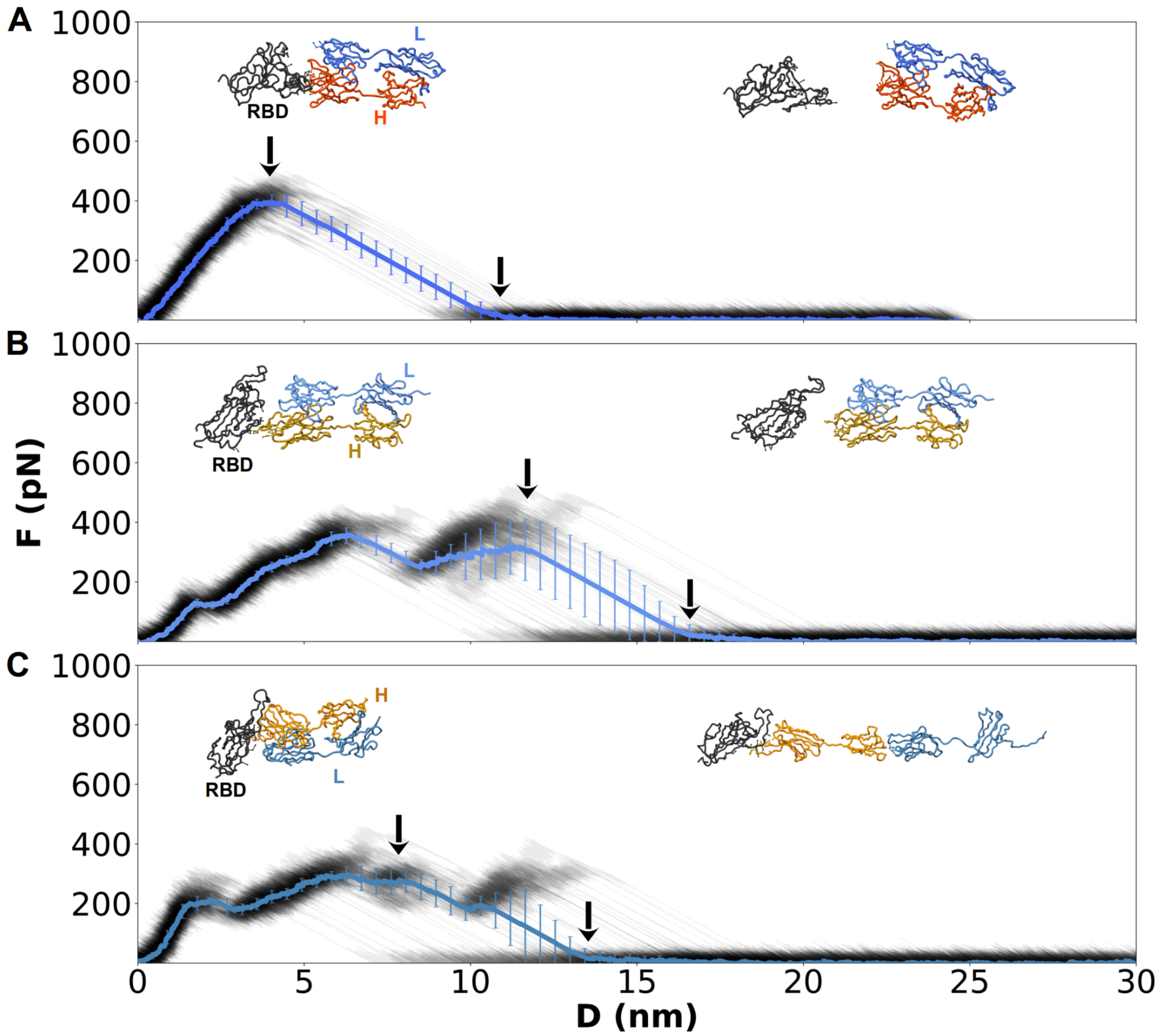


**Figure S7. Mechanical response of Ab/RBD JN.1 complexes under constant-velocity pulling from the L chain.** Same as Figure S4.

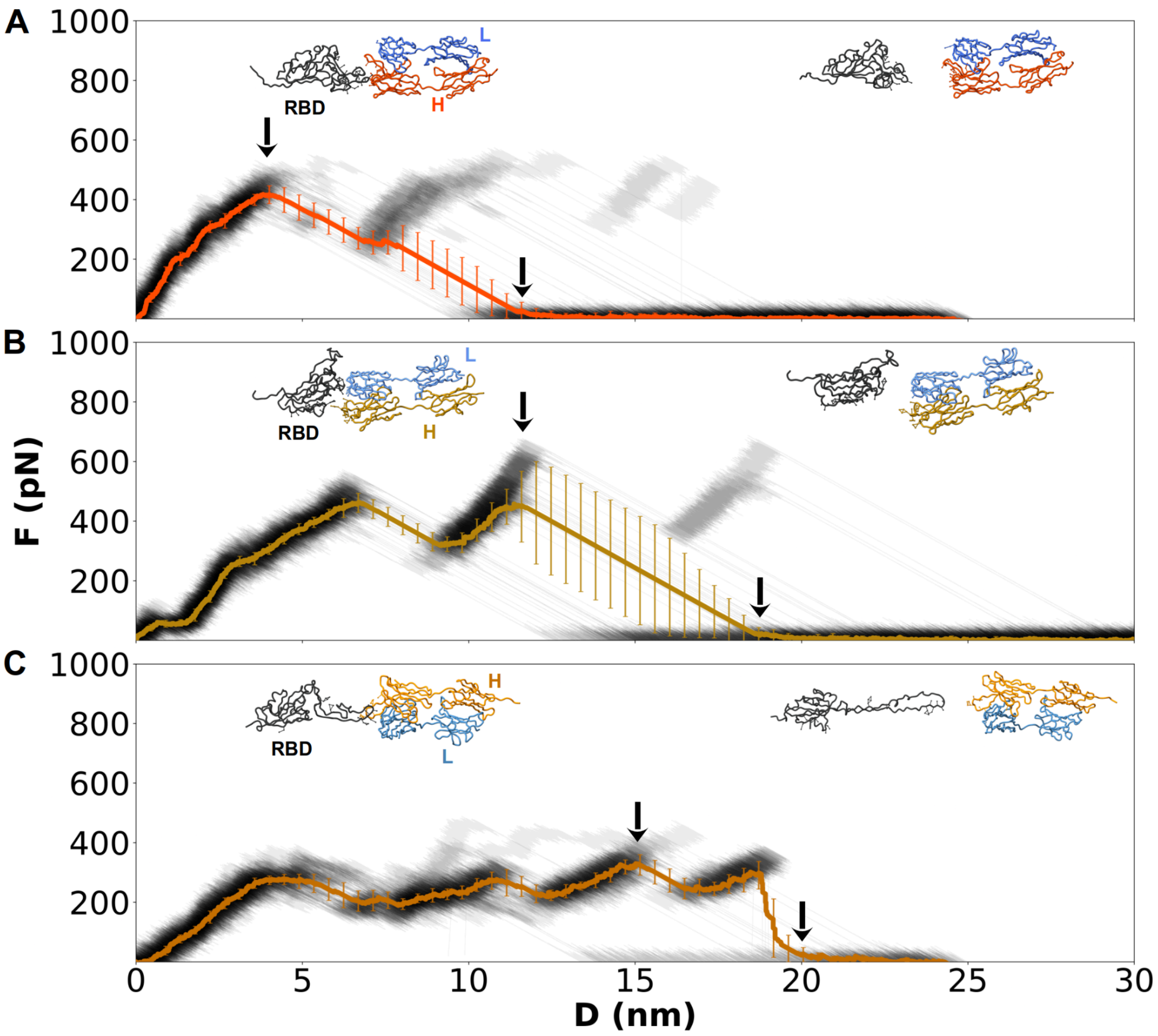


**Figure S8. Mechanical response of Ab/RBD BA.4 complexes under constant-velocity pulling from the H chain.** Same as Figure S4.

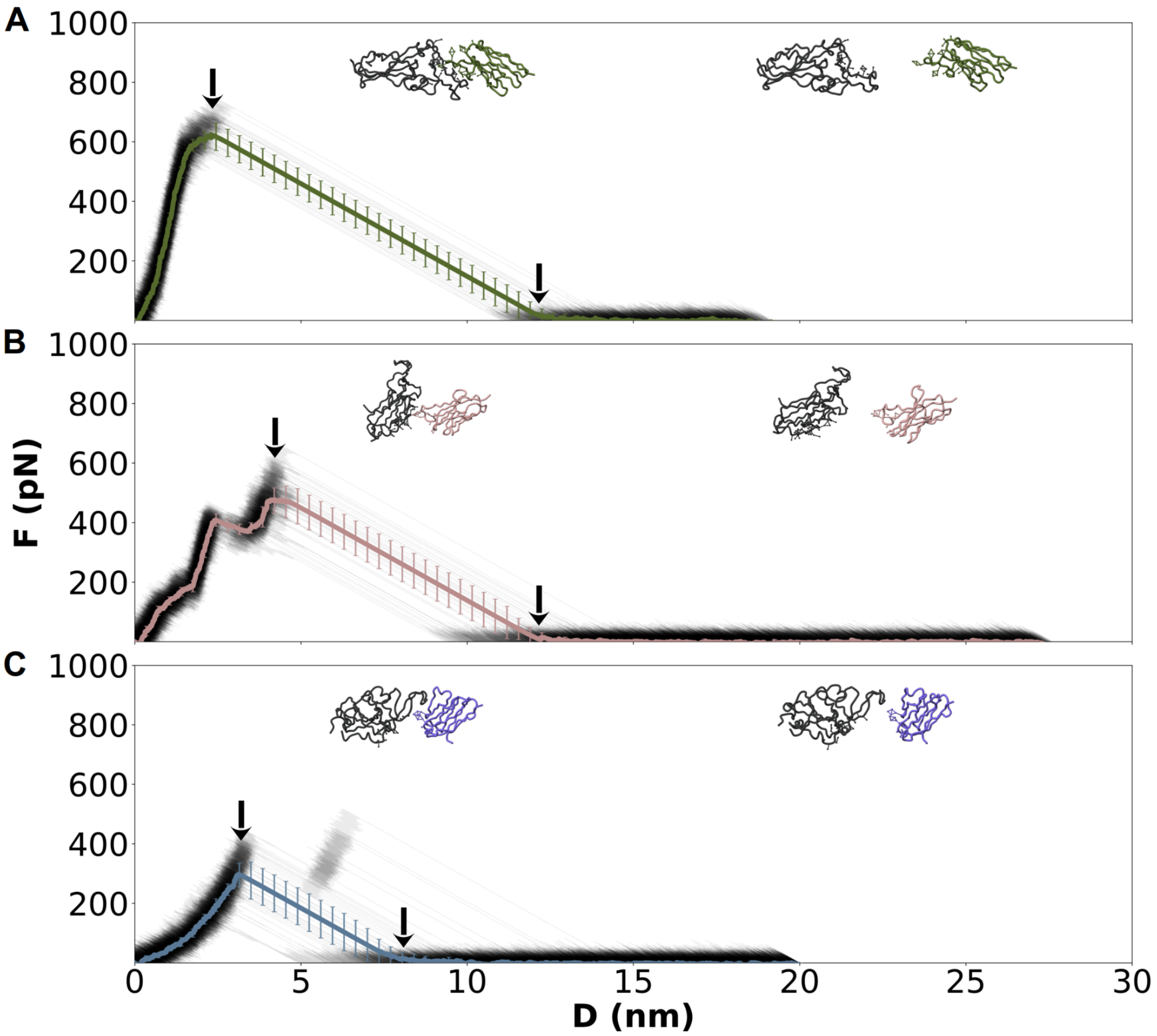


**Figure S9. Mechanical response of Nb/RBD BA.4 complexes under constant-velocity pulling.** Force–displacement profiles are shown for (A) R14, (B) C1, and (C) n313.1 (gray traces, mean in color). Insets display representative structures at the maximum force (Fmax, left arrow) and after dissociation (right arrow). Error bars denote standard deviations across 50 independent trajectories.

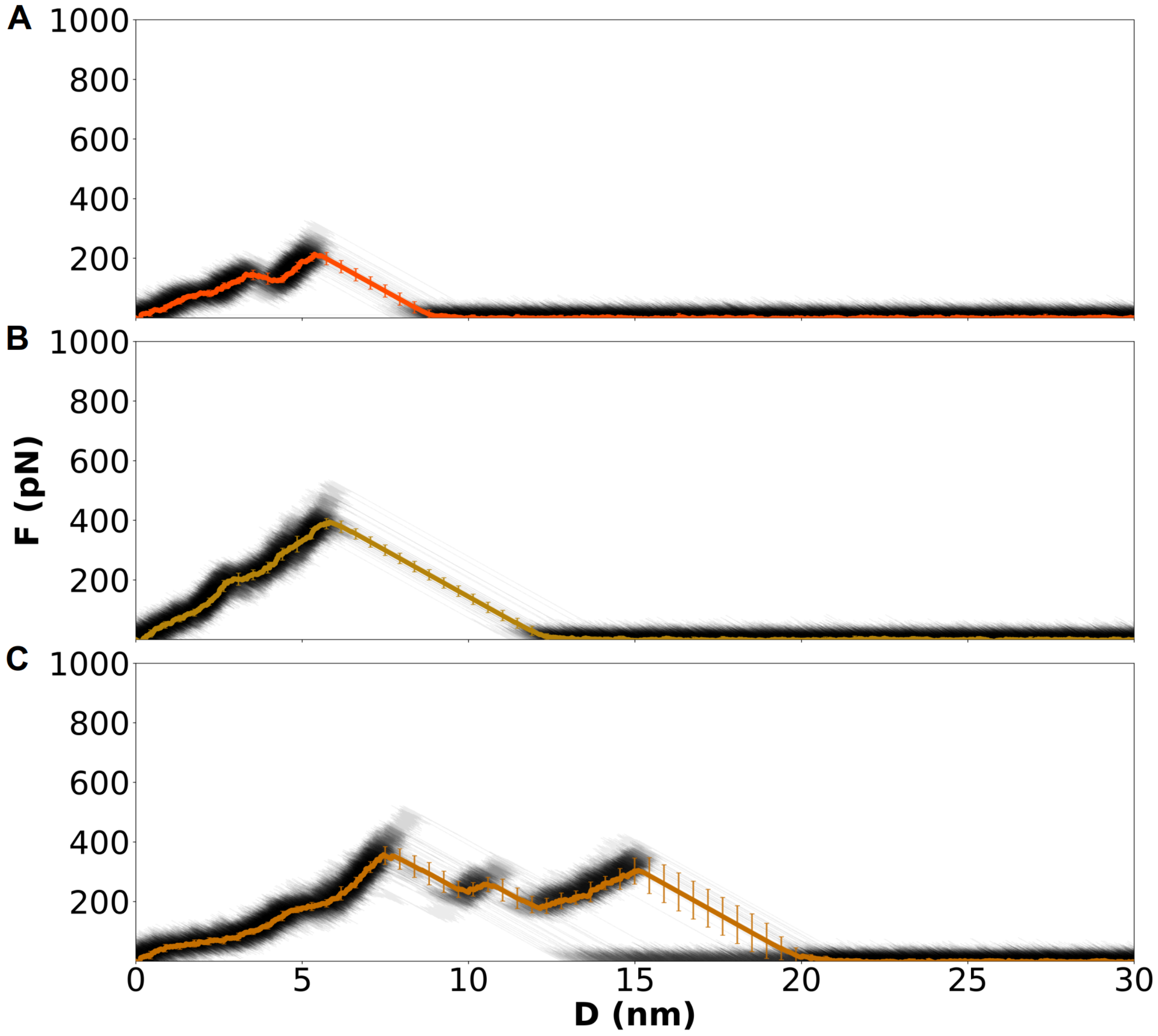


**Figure S10. Mechanical response of H chain/RBD WT complexes under constant-velocity pulling from the H chain.** Same as Figure S4.

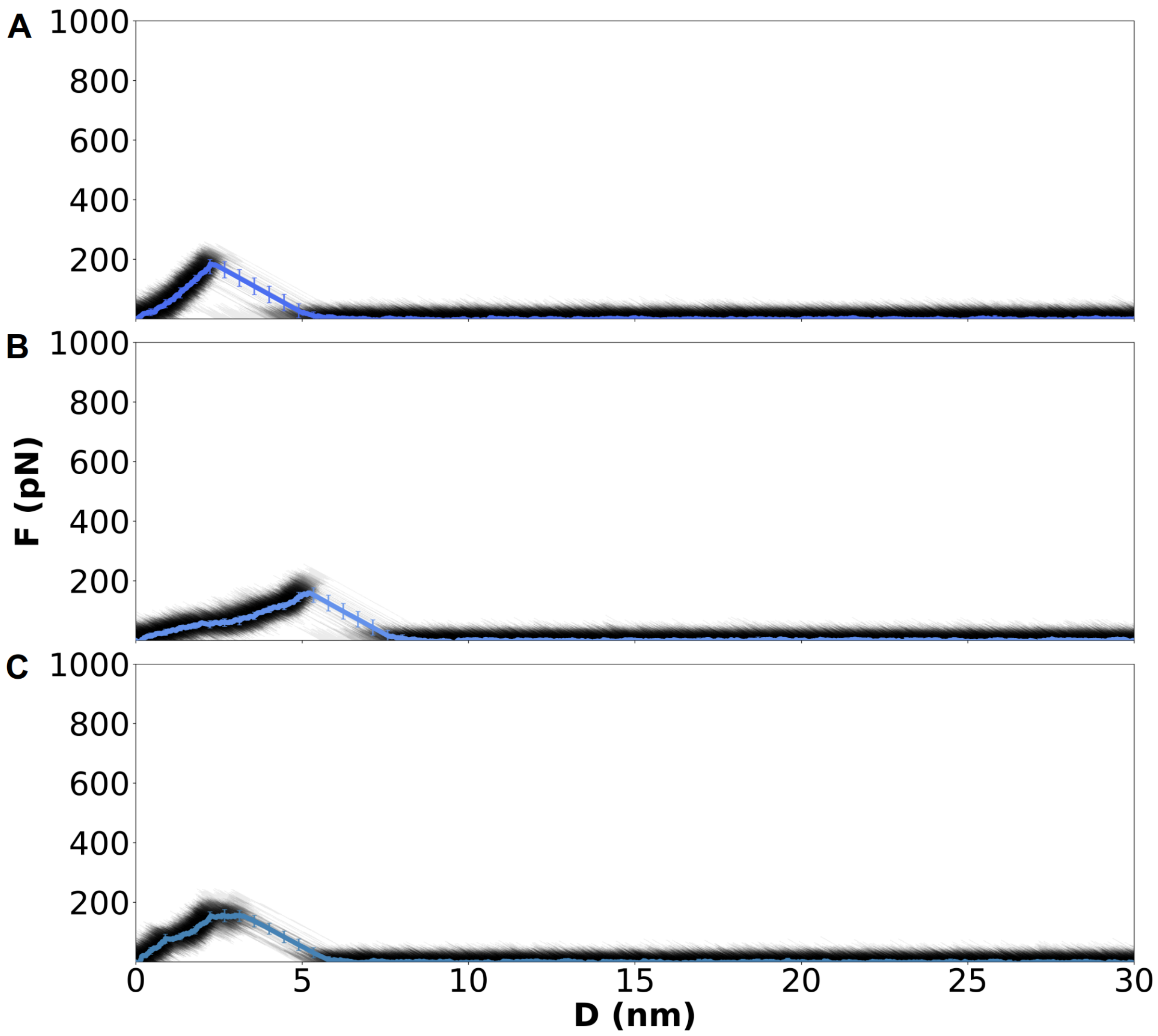


**Figure S11. Mechanical response of L chain/RBD WT complexes under constant-velocity pulling from the L chain.** Same as Figure S4.

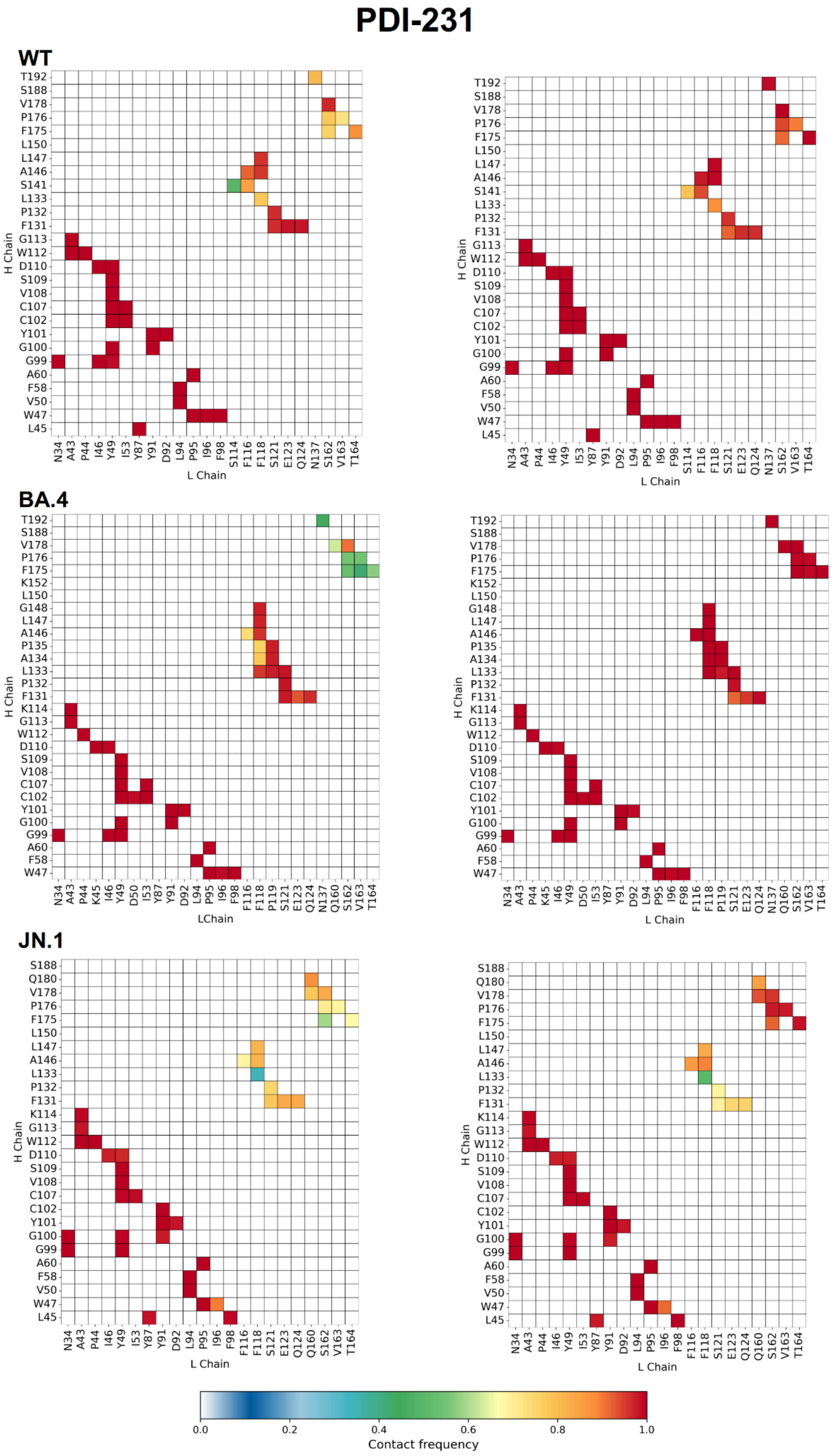


**Figure S12. Interchain contact frequency maps for the PDI-231 antibody under directional pulling.** Contact maps show the frequency of interchain contacts between the heavy (H) and light (L) chains of the PDI-231 antibody in complex with the SARS-CoV-2 RBD for three viral variants: WT (top), BA.4 (middle), and JN.1 (bottom). The left panels correspond to simulations where pulling was applied to the heavy chain, while the right panels correspond to pulling from the light chain. Contacts were calculated using the extended VdW radii method across 50 steered molecular dynamics trajectories. Color intensity reflects contact frequency.

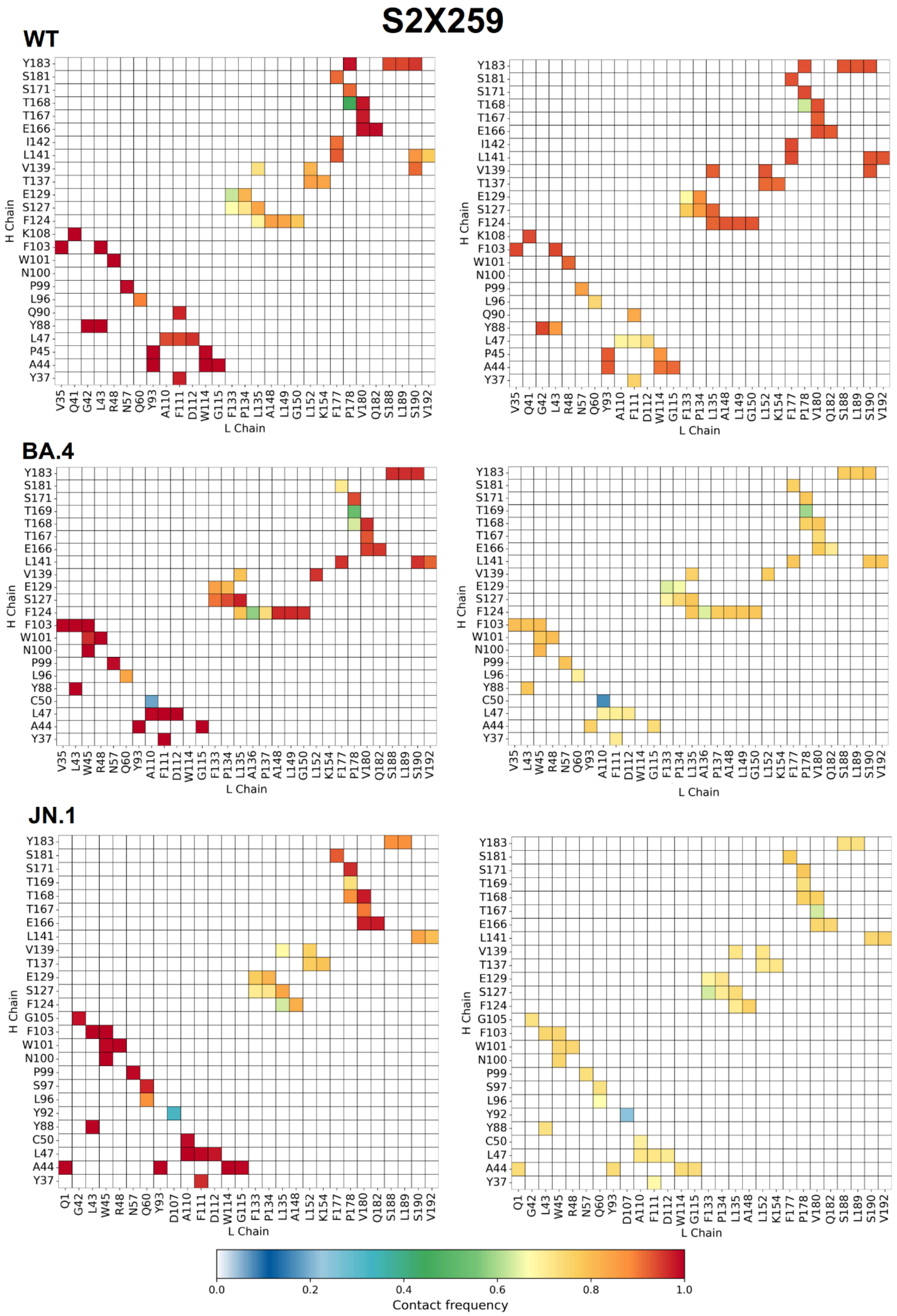


**Figure S13. Interchain contact frequency maps for the S2X259 antibody under directional pulling.** Contact maps show the frequency of interchain contacts between the heavy (H) and light (L) chains of the S2X259 antibody in complex with the SARS-CoV-2 RBD for three viral variants: WT (top), BA.4 (middle), and JN.1 (bottom). The left panels correspond to simulations where pulling was applied to the heavy chain, while the right panels correspond to pulling from the light chain. Contacts were calculated using the extended VdW radii method across 50 steered molecular dynamics trajectories. Color intensity reflects contact frequency.

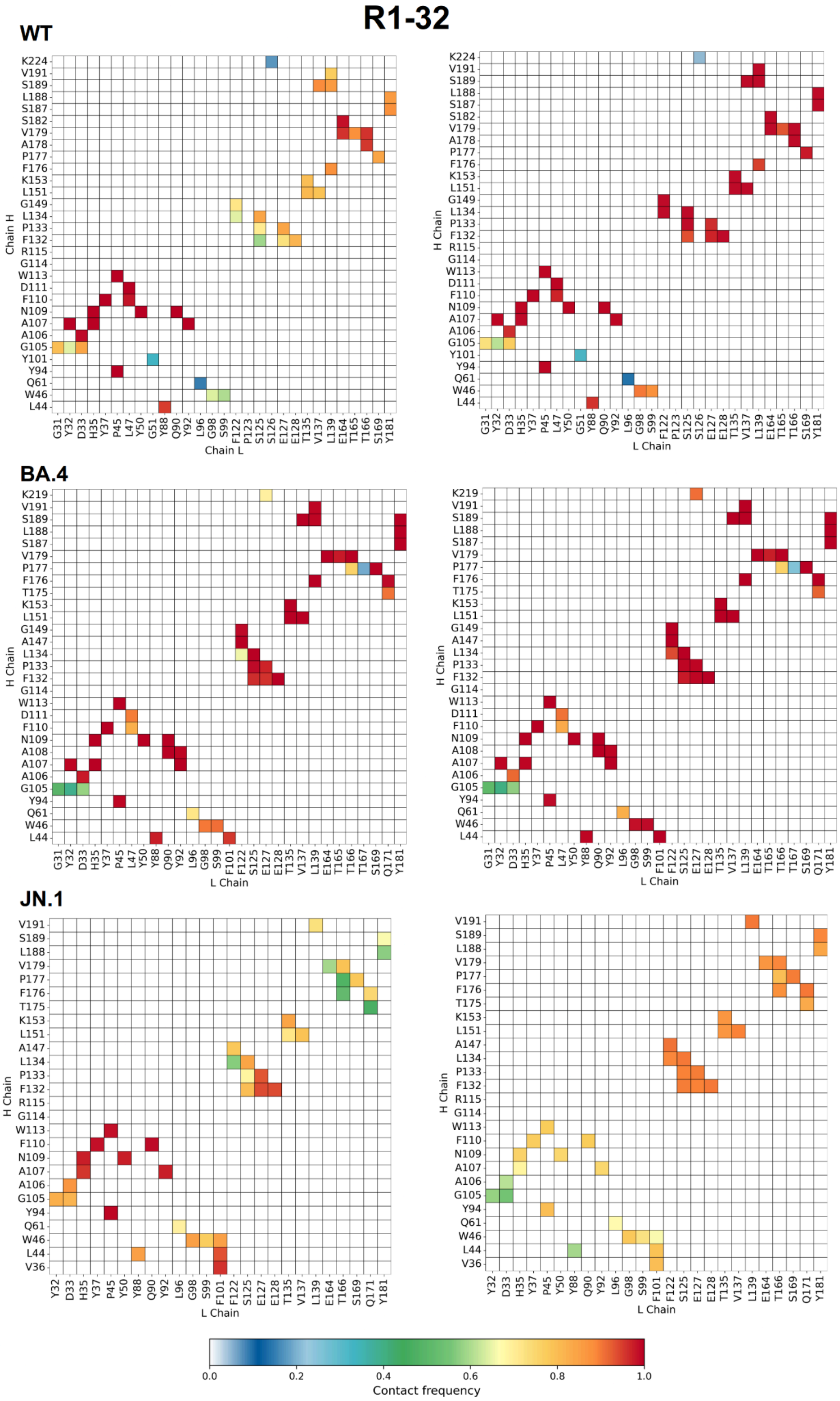


**Figure S14. Interchain contact frequency maps for the R1-32 antibody under directional pulling.** Contact maps show the frequency of interchain contacts between the heavy (H) and light (L) chains of the R1-32 antibody in complex with the SARS-CoV-2 RBD for three viral variants: WT (top), BA.4 (middle), and JN.1 (bottom). The left panels correspond to simulations where pulling was applied to the heavy chain, while the right panels correspond to pulling from the light chain. Contacts were calculated using the extended VdW radii method across 50 steered molecular dynamics trajectories. Color intensity reflects contact frequency.

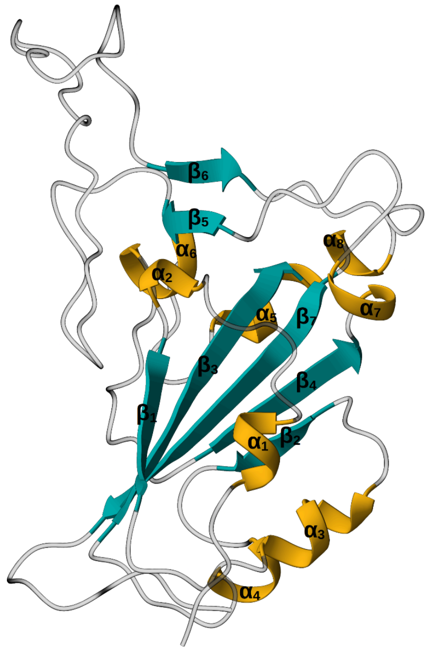


| Helix | Residue range | β-strand | Residue range |
|---|---|---|---|
| $\alpha_1$ | 338-343 | $\beta_1$ | 354-358 |
| $\alpha_2$ | 350-352 | $\beta_2$ | 376-379 |
| $\alpha_3$ | 365-371 | $\beta_3$ | 394-403 |
| $\alpha_4$ | 384-387 | $\beta_4$ | 432-437 |
| $\alpha_5$ | 404-409 | $\beta_5$ | 452-545 |
| $\alpha_6$ | 417-421 | $\beta_6$ | 492-494 |
| $\alpha_7$ | 439-442 | $\beta_7$ | 507-516 |
| $\alpha_8$ | 503-505 | | |

**Figure S15. Secondary structure elements of the SARS-CoV-2 RBD.** The α-helices and β-strands are shown on the RBD cartoon representation, with corresponding residue ranges listed in the table.

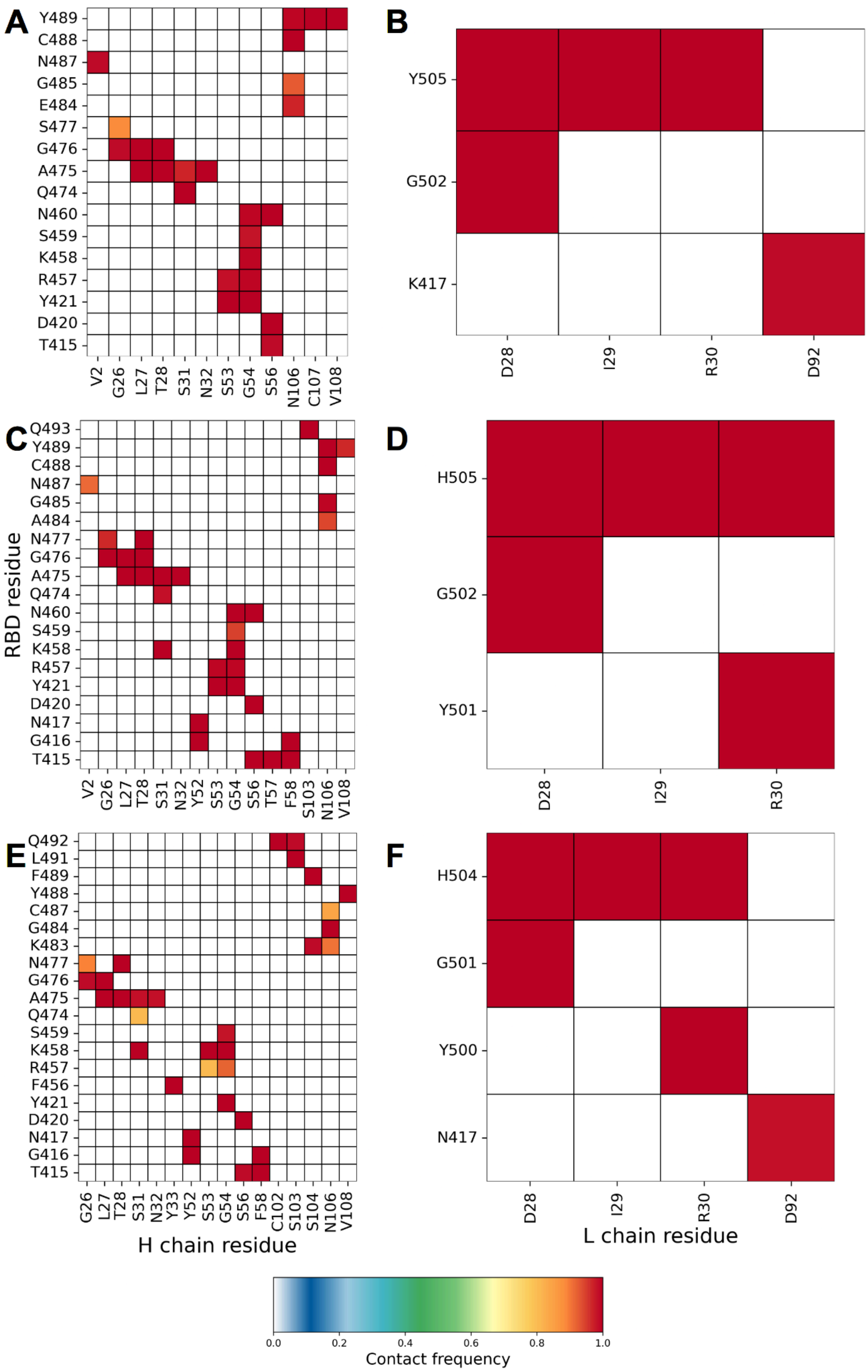


**Figure S16. Interchain contact frequency maps for the RBD/PDI-231 complex from equilibrium simulations.** Contact maps show the frequency of interchain contacts between the RBD and the H or L chains of the PDI-231 Ab. The left panels correspond to contacts with the H chain, and the right panels to contacts with the L chain. Contacts were computed using the extended van der Waals radii method across five independent CG-MD simulations. Color intensity reflects the frequency of each contact. Panels A and B correspond to the WT complex, panels C and D to the BA.4 complex, and panels E and F to the JN.1 complex.

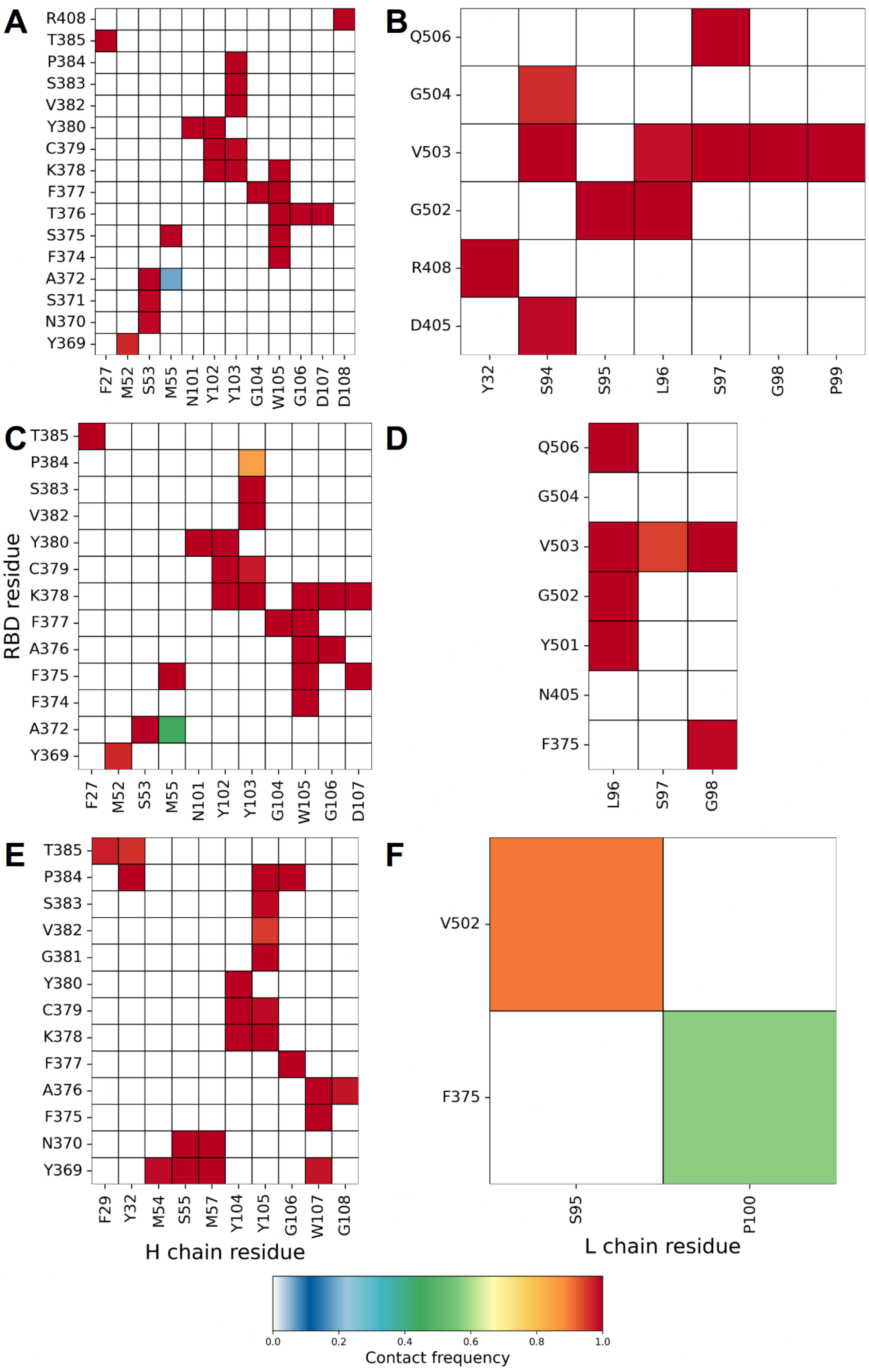


**Figure S17. Interchain contact frequency maps for the RBD/S2X259 complex from equilibrium simulations.** Same as Figure S13.

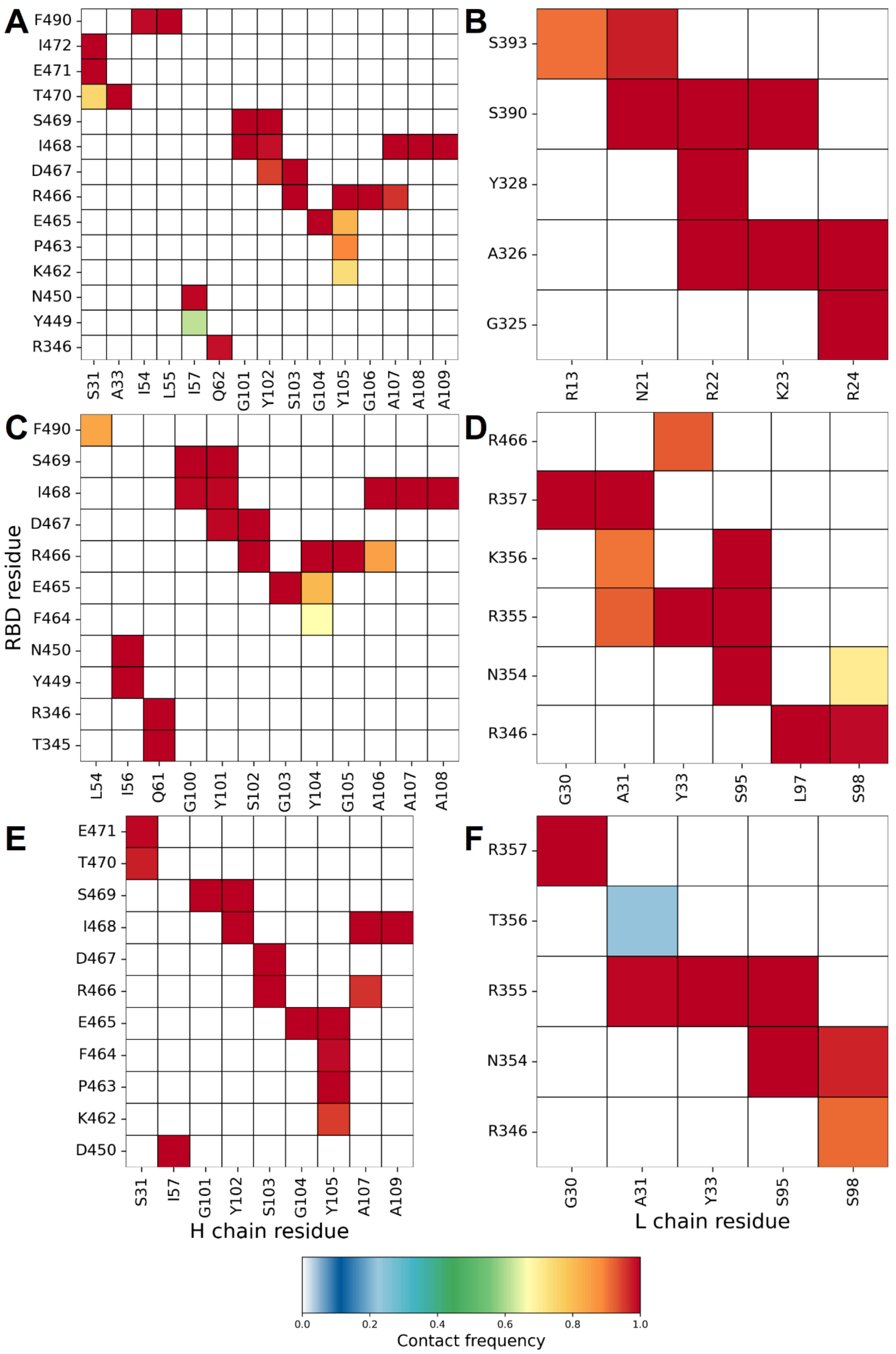


**Figure S18. Interchain contact frequency maps for the RBD/R1-32 complex from equilibrium simulations.** Same as Figure S13.

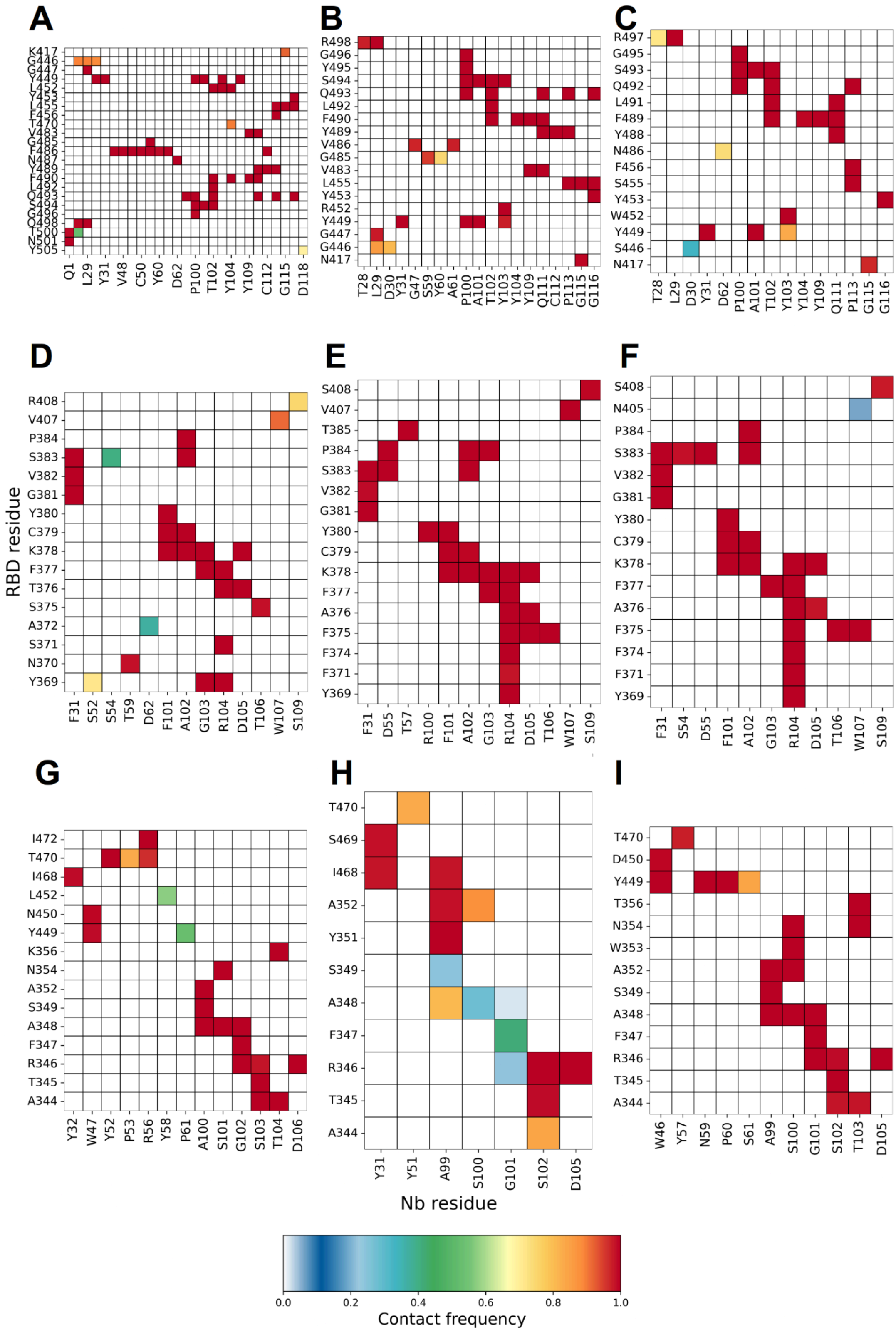


**Figure S19. Interchain contact frequency maps for the RBD/Nb complexes from equilibrium simulations.** Contact maps show the frequency of interchain contacts between the RBD and nanobodies R14, C1, and n3113.1. Contacts were computed using the extended van der Waals radii method across five independent coarse-grained molecular dynamics simulations. Color intensity reflects the frequency of each contact, from rare (blue) to persistent (red). Panels A, D, and G correspond to the WT complex; panels B, E, and H to the BA.4 complex; and panels C, F, and I to the JN.1 complex. Rows correspond to the different nanobodies: the first row to R14 (A–C), the second row to C1 (D–F), and the third row to n3113.1 (G–I).

**Table S1. Structural and binding parameters of the selected nanobodies targeting the SARS-CoV-2 RBD.** For each protein case, the name, type, PDB ID, and RBD residues making contacts with the WT are listed. Experimental dissociation constant (KD) parameters considering different variants are reported. Residues in bold are part of the RBM region.

| Ab | R14 (PDB ID: 7MZN) | | | C1 (PDB ID: 7OAP) | | | n3113.1 (PDB ID: 7VNE) | | |
|---|---|---|---|---|---|---|---|---|---|
| Variant | WT | BA.4 | JN.1 | WT | BA.4 | JN.1 | WT | BA.4 | JN.1 |
| | K417 | N417 | N417 | Y369 | * | * | A344 | * | * |
| | **G446** | * | **S446** | N370 | - | - | T345 | * | * |
| | **G447** | * | - | S371 | - | - | F347 | * | * |
| | **L452** | **R452** | **W452** | A372 | - | * | A348 | * | * |
| | **L455** | * | * | S375 | - | - | A352 | * | * |
| | **F456** | - | - | T376 | - | - | R346 | * | * |
| | **Y449** | * | * | F377 | * | * | S349 | * | * |
| | **Y453** | * | * | C379 | * | * | - | - | **W353** |
| | **F486** | - | **N486** | Y380 | * | * | **N354** | - | * |
| | **F490** | * | * | G381 | * | * | K356 | - | - |
| | **L492** | * | * | V382 | * | * | **Y449** | - | * |
| | **Q493** | * | * | P384 | * | * | **N450** | - | - |
| | **Q498** | **R498** | **R498** | - | - | N405 | **L452** | - | - |
| | **Y489** | * | * | V407 | * | - | **I468** | * | - |
| | **G496** | * | * | R408 | - | - | **T470** | * | - |
| | **S494** | * | * | | | | **I472** | - | - |
| | **T500** | - | - | | | | | | |
| Kd app. (nM) | ND[a,b*31]<br>0.02[c*31]<br>0.03[d*31]<br>43.3[f*3] | | | 0.615[a*32]<br>0.725[b*32]<br>0.648[c*32] | | | 6.4[a**33] | | |

ND refers to no dissociation.

[a]WT
[b]Alpha
[c]Beta
[d]Gamma
[e]Delta
[f]Omicron BA.4/5
*Surface plasmon resonance (SPR)
**Bio-layer interferometry (BLI)

**Table S2. List of protein contacts at the RBD/PDI-231 interface.** Protein-protein interactions were calculated using the OV+rCSU contact map protocol over an MD trajectory of 500 ns. Protein contacts with a frequency of 0.7 were considered in the contact map. The total number of protein contacts is given next to the RBD variant name. Residues from RBD are shown in bold. Hydrophobic, hydrophilic, and ionic contacts are highlighted in green, blue, and red, respectively.

| Chain | WT (31) | BA.4 (38) | JN.1 (39) |
|---|---|---|---|
| H | **A475**-L27 | **A475**-L27 | **A475**-L27 |
| | **A475**-N32 | **A475**-N32 | **A475**-N32 |
| | **A475**-S31 | **A475**-S31 | **A475**-S31 |
| | **A475**-T28 | **A475**-T28 | **A475**-T28 |
| | - | **A484**-N106 | - |
| | **C488**-N106 | **C488**-N106 | **C488**-N106 |
| | **D420**-S56 | **D420**-S56 | **D420**-S56 |
| | **E484**-N106 | - | - |
| | - | - | **F456**-Y33 |
| | - | - | **F489**-S104 |
| | - | **G416**-F58 | **G416**-F58 |
| | - | **G416**-Y52 | **G416**-Y52 |
| | **G476**-G26 | **G476**-G26 | **G476**-G26 |
| | **G476**-L27 | **G476**-L27 | **G476**-L27 |
| | **G476**-T28 | **G476**-T28 | **G476**-T28 |
| | **G485**-N106 | **G485**-N106 | **G485**-N106 |
| | **K458**-G54 | **K458**-G54 | **K458**-G54 |
| | - | **K458**-S31 | **K458**-S31 |
| | - | - | **K458**-S53 |
| | - | - | **K483**-N106 |
| | - | - | **K483**-S104 |
| | - | - | **L492**-S103 |
| | - | **N417**-Y52 | **N417**-Y52 |
| | **N460**-G54 | **N460**-G54 | - |
| | **N460**-S56 | **N460**-S56 | - |
| | - | **N477**-G26 | **N477**-G26 |
| | - | **N477**-T28 | **N477**-T28 |
| | **N487**-V2 | **N487**-V2 | - |
| | **Q474**-S31 | **Q474**-S31 | **Q474**-S31 |
| | - | - | **E493**-C102 |
| | - | **Q493**-S103 | **E493**-S103 |
| | **R457**-G54 | **R457**-G54 | **R457**-G54 |
| | **R457**-S53 | **R457**-S53 | **R457**-S53 |
| | **S459**-G54 | **S459**-G54 | **S459**-G54 |
| | **S477**-G26 | - | - |
| | - | **T415**-F58 | **T415**-F58 |
| | **T415**-S56 | **T415**-S56 | **T415**-S56 |
| | - | **T415**-T57 | - |
| | **Y421**-G54 | **Y421**-G54 | **Y421**-G54 |
| | **Y421**-S53 | **Y421**-S53 | - |
| | **Y489**-C107 | - | - |
| | **Y489**-N106 | **Y489**-N106 | - |
| | **Y489**-V108 | **Y489**-V108 | **Y489**-V108 |
| L | **K417**-D92 | - | **N417**-D92 |
| | - | **Y501**-R30 | **Y501**-R30 |
| | **G502**-D28 | **G502**-D28 | **G502**-D28 |
| | **Y505**-D28 | **H505**-D28 | **H505**-D28 |
| | **Y505**-I29 | **H505**-I29 | **H505**-I29 |
| | **Y505**-R30 | **H505**-R30 | **H505**-R30 |

**Table S3. List of protein contacts at the RBD/S2X259 interface.** Protein-protein interactions were calculated using the OV+rCSU contact map protocol over an MD trajectory of 500 ns. Protein contacts with a frequency of 0.7 were considered in the contact map. The total number of protein contacts is given next to the RBD variant name. Residues from the heavy chain are shown in bold. Hydrophobic, hydrophilic, and ionic contacts are highlighted in green, blue, and red, respectively.

| Chain | WT (36) | BA.4 (37) | JN.1 (25) |
|---|---|---|---|
| H | **A372**-M55 | **A372**-M55 | - |
| | **A372**-S53 | **A372**-S53 | - |
| | - | **A376**-G106 | **A376**-G106 |
| | - | **A376**-W105 | **A376**-W105 |
| | **C379**-Y102 | **C379**-Y102 | **C379**-Y102 |
| | **C379**-Y103 | **C379**-Y103 | **C379**-Y103 |
| | **F374**-W105 | **F374**-W105 | - |
| | - | **F375**-D107 | - |
| | **S375**-M55 | **F375**-M55 | - |
| | **S375**-W105 | **F375**-W105 | **F375**-W105 |
| | **F377**-G104 | **F377**-G104 | **F377**-G104 |
| | **F377**-W105 | **F377**-W105 | - |
| | - | - | **G381**-Y103 |
| | - | **K378**-D107 | - |
| | - | **K378**-G106 | - |
| | **K378**-W105 | **K378**-W105 | - |
| | **K378**-Y102 | **K378**-Y102 | **K378**-Y102 |
| | **K378**-Y103 | **K378**-Y103 | **K378**-Y103 |
| | - | - | **N370**-M55 |
| | **N370**-S53 | - | **N370**-S53 |
| | - | - | **P384**-G104 |
| | **P384**-Y103 | **P384**-Y103 | **P384**-Y103 |
| | - | - | **P384**-Y30 |
| | **R408**-D108 | - | - |
| | **S371**-S53 | - | - |
| | **S383**-Y103 | **S383**-Y103 | **S383**-Y103 |
| | **T376**-D107 | - | - |
| | **T376**-G106 | - | - |
| | **T376**-W105 | - | - |
| | **T385**-F27 | **T385**-F27 | **T385**-F27 |
| | - | - | **T385**-Y30 |
| | **V382**-Y103 | **V382**-Y103 | **V382**-Y103 |
| | **Y369**-M52 | - | **Y369**-M52 |
| | - | - | **Y369**-M55 |
| | - | - | **Y369**-S53 |
| | - | - | **Y369**-W105 |
| | **Y380**-N101 | **Y380**-N101 | - |
| | **Y380**-Y102 | **Y380**-Y102 | **Y380**-Y102 |
| L | **D405**-S95 | - | - |
| | - | **F375**-P100 | **F375**-P100 |
| | **G502**-L97 | **G502**-L97 | - |
| | - | **G502**-S95 | - |
| | **G502**-S96 | - | - |
| | - | **G502**-S98 | - |
| | **G504**-S95 | **G504**-S95 | - |
| | - | **N405**-Y33 | - |
| | **Q506**-S98 | **Q506**-S98 | - |
| | **R408**-Y33 | - | - |
| | **V503**-G99 | **V503**-G99 | - |
| | **V503**-L97 | **V503**-L97 | - |
| | **V503**-P100 | **V503**-P100 | - |
| | **V503**-S95 | **V503**-S95 | **V503**-S95 |
| | **V503**-S98 | **V503**-S98 | - |
| | - | **Y501**-S98 | - |

**Table S4. List of protein contacts at the RBD/R1-32 interface.** Protein-protein interactions were calculated using the OV+rCSU contact map protocol over an MD trajectory of 500 ns. Protein contacts with a frequency of 0.7 were considered in the contact map. The total number of protein contacts is given next to the RBD variant name. Residues from the heavy chain are shown in bold. Hydrophobic, hydrophilic, and ionic contacts are highlighted in green, blue, and red, respectively.

| Chain | WT (36) | BA.4 (33) | JN.1 (24) |
|---|---|---|---|
| | - | - | **D450**-I57 |
| | - | **D467**-S103 | **D467**-S103 |
| | - | **D467**-G101 | - |
| | **D467**-Y102 | - | - |
| | - | **E465**-S103 | - |
| | - | **E465**-G104 | **E465**-G104 |
| | **E465**-Y105 | - | **E465**-Y105 |
| | **E471**-S31 | - | **E471**-S31 |
| | - | **F464**-G104 | - |
| | - | - | **F464**-Y105 |
| | **F490**-I54 | **F490**-I54 | - |
| | **F490**-L55 | - | - |
| | **E465**-G104 | - | - |
| | - | **I468**-G106 | - |
| | **I468**-A107 | **I468**-A107 | **I468**-A107 |
| | **I468**-A108 | **I468**-A108 | - |
| | **I468**-A109 | - | **I468**-A109 |
| | - | **I468**-N100 | - |
| | **I468**-G101 | **I468**-G101 | - |
| | - | - | **I468**-Y102 |
| | **I57**-N450 | | - |
| H | **K462**-Y105 | - | **K462**-Y105 |
| | - | **N450**-G56 | - |
| | **P463**-Y105 | - | **P463**-Y105 |
| | - | **R346**-A61 | - |
| | **R346**-Q62 | - | - |
| | - | **R466**-G106 | - |
| | **R466**-A107 | - | **R466**-A107 |
| | - | **R466**-Y105 | - |
| | **R466**-G106 | - | - |
| | - | **R466**-Y102 | - |
| | - | - | **R466**-S103 |
| | - | **R466**-G104 | - |
| | **R466**-Y105 | - | - |
| | **S103**-D467 | - | - |
| | **S103**-R466 | - | - |
| | **I472**-S31 | - | - |
| | - | **S469**-N100 | - |
| | **S469**-G101 | **S469**-G101 | **S469**-G101 |
| | - | - | **S469**-Y102 |
| | - | **T345**-A61 | - |
| | **T470**-A33 | - | - |
| | **T470**-S31 | - | **T470**-S31 |
| | **I468**-Y102 | - | - |
| | **S469-Y102** | - | - |
| | - | **Y449**-G56 | - |
| | **Y449**-I57 | - | - |
| | **A326**-K23 | - | - |
| | **A326**-R22 | - | - |
| | **A326**-R24 | - | - |
| | **G325**-R24 | - | - |
| | - | **K356**-A31 | - |
| | - | **K356**-S95 | - |
| | - | **N354**-S95 | **N354**-S95 |
| | - | **N354**-S98 | **N354**-S98 |
| | - | **R346**-L97 | - |
| | - | **R346**-S98 | **R346**-S98 |
| | - | **R355**-A31 | **R355**-A31 |
| L | - | **R355**-S95 | **R355**-S95 |
| | - | **R355**-Y33 | **R355**-Y33 |
| | - | **R357**-A31 | - |
| | - | **R357**-G30 | **R357**-G30 |
| | - | **R466**-Y33 | - |
| | **S390**-K23 | - | - |

| | | | |
|---|---|---|---|
| | **S390**-N21<br>**S390**-R22<br>**S393**-N21<br>**S393**-R13<br>-<br>**Y328**-R22 | -<br>-<br>-<br>-<br>-<br>- | -<br>-<br>-<br>-<br>**T356**-A31<br>- |

**Table S5. List of protein contacts at the RBD/R14 interface.** Protein-protein interactions were calculated using the OV+rCSU contact map protocol over an MD trajectory of 500 ns. Protein contacts with a frequency of 0.7 were considered in the contact map. The total number of protein contacts is given next to the RBD variant name. Residues from the heavy chain are shown in bold. Hydrophobic, hydrophilic, and ionic contacts are highlighted in green, blue, and red, respectively.

| WT (41) | BA.4 (40) | JN.1 (26) |
|---|---|---|
| **F456**-P113 | - | **F456**-P113 |
| **F486**-A61 | - | - |
| **F486**-C50 | - | - |
| **F486**-G47 | - | - |
| **F486**-S49 | - | - |
| **F486**-S59 | - | - |
| **F486**-V48 | - | - |
| **F486**-Y60 | - | - |
| **F490**-Q111 | **F490**-Q111 | **F490**-Q111 |
| **F490**-T102 | **F490**-T102 | **F490**-T102 |
| **F490**-Y104 | **F490**-Y104 | **F490**-Y104 |
| **F490**-Y109 | **F490**-Y109 | **F490**-Y109 |
| **G446**-D30 | **G446**-D30 | **S446**-D30 |
| **G446**-L29 | **G446**-L29 | - |
| **G446**-T28 | - | - |
| **G447**-L29 | **G447**-L29 | - |
| - | **G485**-S59 | - |
| - | **G485**-Y60 | - |
| **G496**-P100 | **G496**-P100 | **G496**-P100 |
| **K417**-G115 | **N417**-G115 | **N417**-G115 |
| **L452**-Y103 | **R452**-Y103 | **W452**-Y103 |
| **L455**-G115 | **L455**-G115 | - |
| **L455**-G116 | **L455**-G116 | - |
| **L455**-P113 | **L455**-P113 | **S455**-P113 |
| - | - | **L492**-Q111 |
| - | - | **N486**-D62 |
| **L492**-T102 | **L492**-T102 | **L492**-T102 |
| **Q493**-G116 | **Q493**-G116 | - |
| **Q493**-P100 | **Q493**-P100 | **Q493**-P100 |
| - | **Q493**-P113 | **Q493**-P113 |
| - | **Q493**-Q111 | - |
| **Q493**-T102 | **Q493**-T102 | **Q493**-T102 |
| **Q498**-L29 | **R498**-L29 | **R498**-L29 |
| **Q498**-T28 | **R498**-T28 | **R498**-T28 |
| **S494**-A101 | **S494**-A101 | **S494**-A101 |
| **S494**-P100 | **S494**-P100 | **S494**-P100 |
| **S494**-T102 | **S494**-T102 | **S494**-T102 |
| - | **S494**-Y103 | - |
| **T500**-T28 | - | - |
| - | **V483**-Q111 | - |
| - | **V483**-Y109 | - |
| - | **V486**-A61 | - |
| - | **V486**-G47 | - |
| **Y449**-A101 | **Y449**-A101 | **Y449**-A101 |
| **Y449**-D30 | - | - |
| **Y449**-P100 | **Y449**-P100 | - |
| **Y449**-Y103 | **Y449**-Y103 | **Y449**-Y103 |
| **Y449**-Y31 | **Y449**-Y31 | **Y449**-Y31 |
| **Y453**-G116 | **Y453**-G116 | **Y453**-G116 |
| - | - | |
| **Y489**-C112 | **Y489**-C112 | - |
| **Y489**-P113 | **Y489**-P113 | - |
| **Y489**-Q111 | **Y489**-Q111 | **Y489**-Q111 |
| - | **Y495**-P100 | - |

**Table S6. List of protein contacts at the RBD/C1 interface.** Protein-protein interactions were calculated using the OV+rCSU contact map protocol over an MD trajectory of 500 ns. Protein contacts with a frequency of 0.7 were considered in the contact map. The total number of protein contacts is given next to the RBD variant name. Residues from the heavy chain are shown in bold. Hydrophobic, hydrophilic, and ionic contacts are highlighted in green, blue, and red, respectively.

| WT (26) | BA.4 (30) | JN.1 (34) |
|---|---|---|
| **A372**-D62 | - | - |
| - | - | **A372**-R104 |
| - | **A376**-D105 | **A376**-D105 |
| - | **A376**-R104 | **A376**-R104 |
| **C379**-A102 | **C379**-A102 | **C379**-A102 |
| **C379**-F101 | **C379**-F101 | **C379**-F101 |
| - | - | **C379**-F31 |
| - | **F371**-R104 | **F371**-R104 |
| - | **F374**-R104 | **F374**-R104 |
| - | **F375**-D105 | - |
| - | **F375**-R104 | **F375**-R104 |
| - | **F375**-T106 | **F375**-T106 |
| - | - | **F375**-W107 |
| **F377**-G103 | **F377**-G103 | **F377**-G103 |
| **F377**-R104 | **F377**-R104 | **F377**-R104 |
| **G381**-F31 | **G381**-F31 | **G381**-F31 |
| **K378**-A102 | **K378**-A102 | **K378**-A102 |
| **K378**-D105 | **K378**-D105 | **K378**-D105 |
| **K378**-F101 | **K378**-F101 | **K378**-F101 |
| **K378**-G103 | **K378**-G103 | - |
| - | **K378**-R104 | **K378**-R104 |
| - | - | **K378**-S110 |
| **N370**-T59 | - | - |
| - | - | **N405**-W107 |
| **P384**-A102 | **P384**-A102 | **P384**-A102 |
| - | **P384**-D55 | - |
| - | **P384**-G103 | - |
| **R408**-S109 | - | - |
| **S371**-R104 | - | - |
| **S375**-T106 | - | - |
| **S383**-A102 | **S383**-A102 | **S383**-A102 |
| - | **S383**-D55 | **S383**-D55 |
| **S383**-F31 | **S383**-F31 | **S383**-F31 |
| **S383**-S54 | - | **S383**-S54 |
| - | **S408**-S109 | **S408**-S109 |
| **T376**-D105 | - | - |
| **T376**-R104 | - | - |
| - | - | **T385**-D55 |
| - | **T385**-T57 | **T385**-T57 |
| **V382**-F31 | **V382**-F31 | **V382**-F31 |
| **V407**-W107 | **V407**-W107 | - |
| **Y369**-G103 | - | **Y369**-G103 |
| **Y369**-R104 | **Y369**-R104 | **Y369**-R104 |
| **Y369**-S52 | - | - |
| - | - | **Y369**-T57 |
| **Y380**-F101 | **Y380**-F101 | **Y380**-F101 |
| - | - | **Y380**-F31 |
| - | **Y380**-R100 | - |

**Table S7. List of protein contacts at the RBD/n3113.1 interface.** Protein-protein interactions were calculated using the OV+rCSU contact map protocol over an MD trajectory of 500 ns. Protein contacts with a frequency of 0.7 were considered in the contact map. The total number of protein contacts is given next to the RBD variant name. Residues from the heavy chain are shown in bold. Hydrophobic, hydrophilic, and ionic contacts are highlighted in green, blue, and red, respectively.

| n3113.1 (PDB ID: 7VNE) | | |
|---|---|---|
| WT (23) | BA.4 (17) | JN.1 (23) |
| **A344**-S103 | **A344**-S103 | **A344**-S1023 |
| **A344**-T104 | - | **A344**-T104 |
| **A348**-A100 | **A348**-A100 | **A348**-A100 |
| **A348**-G102 | **A348**-G102 | **A348**-G102 |
| **A348**-S101 | **A348**-S101 | **A348**-S101 |
| **A352**-A100 | **A352**-A100 | **A352**-A100 |
| - | **A352**-S101 | **A352**-S101 |
| - | - | **D450**-W47 |
| **F347**-G102 | **F347**-G102 | **F347**-G102 |
| - | - | - |
| - | **I468**-A100 | - |
| **I468**-Y32 | **I468**-Y32 | - |
| **I472**-R56 | - | - |
| **K356**-T104 | - | - |
| **L452**-Y58 | - | - |
| **N354**-S101 | - | **N354**-S101 |
| **N450**-W47 | - | - |
| - | - | **N354**-T104 |
| **R346**-D106 | **R346**-D106 | **R346**-D106 |
| **R346**-G102 | **R346**-G102 | **R346**-G102 |
| **R346**-S103 | **R346**-S103 | **R346**-S103 |
| **S349**-A100 | **S349**-A100 | **S349**-A100 |
| - | **S469**-Y32 | - |
| **T345**-S103 | **T345**-S103 | **T345**-S103 |
| **T470**-P53 | - | - |
| **T470**-R56 | - | - |
| - | - | **T356**-T104 |
| **T470**-Y52 | **T470**-Y52 | - |
| - | - | - |
| - | **Y351**-A100 | - |
| - | | **T470**-Y58 |
| - | - | **W353**-S101 |
| - | - | **Y449**-N60 |
| **Y449**-P61 | - | **Y449**-P61 |
| - | - | **Y449**-S62 |
| **Y449**-W47 | - | **Y449**-W47 |

**Table S8. Interchain contacts at the PDI-231 Ab interface.** Contacts were identified using the OV+rCSU contact map protocol applied to a 500 ns molecular dynamics trajectory. Only interactions with a frequency of 0.7 or higher were included. Each contact is denoted as a residue pair, where the first residue belongs to the heavy chain and the second to the light chain. The total number of inter-chain contacts is reported for each SARS-CoV-2 RBD variant. Hydrophobic, hydrophilic, and ionic contacts are highlighted in green, blue, and red, respectively.

| WT (49) | BA.4 (55) | JN.1 (45) |
|---|---|---|
| - | A134-F118 | - |
| - | A134-P119 | - |
| A146-F116 | A146-F116 | A146-F116 |
| A146-F118 | A146-F118 | A146-F118 |
| A60-P95 | A60-P95 | A60-P95 |
| C102-D50 | C102-D50 | C102-D50 |
| C102-I53 | C102-I53 | - |
| C102-Y49 | C102-Y49 | - |
| - | - | C102-Y91 |
| C107-I53 | C107-I53 | C107-I53 |
| C107-Y49 | C107-Y49 | C107-Y49 |
| D110-I46 | D110-I46 | D110-I46 |
| D110-K45 | D110-K45 | - |
| D110-Y49 | - | D110-Y49 |
| F131-E123 | F131-E123 | F131-E123 |
| F131-Q124 | F131-Q124 | F131-Q124 |
| F131-S121 | F131-S121 | F131-S121 |
| F175-S162 | F175-S162 | F175-S162 |
| F175-T164 | F175-T164 | F175-T164 |
| - | F175-V163 | - |
| F58-L94 | F58-L94 | F58-L94 |
| - | - | G100-N34 |
| G100-Y49 | G100-Y49 | G100-Y49 |
| G100-Y91 | G100-Y91 | G100-Y91 |
| G113-A43 | G113-A43 | G113-A43 |
| - | G148-F118 | - |
| G99-I46 | G99-I46 | - |
| G99-N34 | G99-N34 | G99-N34 |
| G99-Y49 | G99-Y49 | G99-Y49 |
| K114-A43 | K114-A43 | K114-A43 |
| - | K152-S131 | - |
| L133-F118 | L133-F118 | L133-F118 |
| - | L133-P119 | - |
| - | L133-S121 | - |
| L147-F118 | L147-F118 | L147-F118 |
| L150-S131 | L150-S131 | L150-S131 |
| - | L150-V133 | - |
| - | L45-F98 | L45-F98 |
| L45-Y87 | L45-Y87 | L45-Y87 |
| P132-S121 | P132-S121 | P132-S121 |
| - | P135-F118 | - |
| - | P135-P119 | - |
| P176-S162 | P176-S162 | P176-S162 |
| P176-V163 | P176-V163 | P176-V163 |
| - | - | Q180-Q160 |
| S109-Y49 | S109-Y49 | S109-Y49 |
| S141-F116 | - | - |
| S141-S114 | - | - |
| S188-S176 | S188-S176 | S188-S176 |
| T192-N137 | T192-N137 | - |
| V108-Y49 | V108-Y49 | V108-Y49 |
| - | V178-Q160 | V178-Q160 |
| V178-S162 | V178-S162 | V178-S162 |
| V50-L94 | - | V50-L94 |
| W112-A43 | - | W112-A43 |
| W112-K45 | - | - |
| W112-P44 | W112-P44 | W112-P44 |
| W47-F98 | W47-F98 | - |
| W47-I96 | W47-I96 | W47-I96 |
| W47-P95 | W47-P95 | W47-P95 |

| | | |
|---|---|---|
| Y101-D92<br>Y101-Y91<br>Y94-A43<br>Y94-P44 | Y101-D92<br>Y101-Y91<br>Y94-A43<br>Y94-P44 | Y101-D92<br>Y101-Y91<br>Y94-A43<br>Y94-P44 |

**Table S9. Interchain contacts at the S2X259 antibody interface.** Contacts were identified using the OV+rCSU contact map protocol applied to a 500 ns molecular dynamics trajectory. Only interactions with a frequency of 0.7 or higher were included. Each contact is denoted as a residue pair, where the first residue belongs to the heavy chain and the second to the light chain. The total number of inter-chain contacts is reported for each SARS-CoV-2 RBD variant. Hydrophobic, hydrophilic, and ionic contacts are highlighted in green, blue, and red, respectively.

| WT (46) | BA.4 (49) | JN.1 (46) |
|---|---|---|
| - | A110-C50 | A110-C50 |
| A110-L47 | A110-L47 | A110-L47 |
| - | A136-F124 | - |
| A148-F124 | A148-F124 | A148-F124 |
| - | - | D107-Y92 |
| D112-L47 | D112-L47 | D112-L47 |
| - | E44-F103 | - |
| F111-L47 | F111-L47 | F111-L47 |
| F111-Q90 | - | - |
| F111-Y37 | F111-Y37 | F111-Y37 |
| F133-E129 | F133-E129 | F133-E129 |
| F133-S127 | F133-S127 | F133-S127 |
| F177-I142 | - | - |
| F177-L141 | F177-L141 | - |
| F177-S181 | F177-S181 | F177-S181 |
| G115-A44 | G115-A44 | G115-A44 |
| G150-F124 | G150-F124 | - |
| G42-G105 | - | G42-G105 |
| K154-T137 | K154-T137 | K154-T137 |
| L135-F124 | L135-F124 | L135-F124 |
| L135-S127 | L135-S127 | L135-S127 |
| L135-V139 | L135-V139 | L135-V139 |
| L149-F124 | L149-F124 | - |
| L152-T137 | L152-T137 | L152-T137 |
| L152-V139 | L152-V139 | L152-V139 |
| L189-Y183 | L189-Y183 | L189-Y183 |
| L43-F103 | L43-F103 | L43-F103 |
| L43-Y88 | L43-Y88 | L43-Y88 |
| N57-P99 | N57-P99 | N57-P99 |
| P134-E129 | P134-E129 | P134-E129 |
| P134-S127 | P134-S127 | P134-S127 |
| P137-F124 | P137-F124 | - |
| P178-S171 | P178-S171 | P178-S171 |
| P178-T168 | P178-T168 | P178-T168 |
| - | P178-T169 | P178-T169 |
| P178-Y183 | - | - |
| - | - | Q1-A44 |
| Q182-E166 | Q182-E166 | Q182-E166 |
| Q41-K108 | - | - |
| Q60-L96 | Q60-L96 | Q60-L96 |
| - | - | Q60-S97 |
| R48-W101 | R48-W101 | R48-W101 |
| - | S183-E166 | S183-E166 |
| S188-Y183 | S188-Y183 | S188-Y183 |
| S190-T141 | S190-T141 | S190-T141 |
| S190-V139 | - | - |
| S190-Y183 | S190-Y183 | - |
| V180-E166 | V180-E166 | V180-E166 |
| V180-T167 | V180-T167 | V180-T167 |
| V180-T168 | V180-T168 | V180-T168 |
| V192-L141 | V192-L141 | V192-L141 |
| V35-F103 | V35-F103 | - |
| W114-A44 | - | W114-A44 |
| W114-P45 | W114-P45 | W114-P45 |
| W45-F103 | W45-F103 | W45-F103 |
| W45-N100 | W45-N100 | W45-N100 |
| W45-W101 | W45-W101 | W45-W101 |
| Y93-A44 | Y93-A44 | Y93-A44 |
| Y93-P45 | Y93-P45 | Y93-P45 |

**Table S10. Interchain contacts at the R1-32 antibody interface.** Contacts were identified using the OV+rCSU contact map protocol applied to a 500 ns molecular dynamics trajectory. Only interactions with a frequency of 0.7 or higher were included. Each contact is denoted as a residue pair, where the first residue belongs to the heavy chain and the second to the light chain. The total number of inter-chain contacts is reported for each SARS-CoV-2 RBD variant. Hydrophobic, hydrophilic, and ionic contacts are highlighted in green, blue, and red, respectively.

| WT (52) | BA.4 (54) | JN.1 (45) |
|---|---|---|
| A106-D33 | A106-D33 | A106-D33 |
| A107-H35 | A107-H35 | A107-H35 |
| A107-Y32 | A107-Y32 | - |
| A107-Y92 | A107-Y92 | A107-Y92 |
| - | A108-Q90 | - |
| - | A108-Y92 | - |
| - | A135-F122 | A135-F122 |
| A135-P123 | - | - |
| - | A147-F122 | A147-F122 |
| A178-T166 | - | - |
| D111-L47 | D111-L47 | - |
| F110-L47 | F110-L47 | - |
| - | - | F110-Q90 |
| F110-Y37 | F110-Y37 | F110-Y37 |
| F132-E127 | F132-E127 | F132-E127 |
| F132-E128 | F132-E128 | F132-E128 |
| F132-S125 | F132-S125 | F132-S125 |
| F176-L139 | F176-L139 | - |
| - | F176-Q171 | F176-Q171 |
| - | - | F176-T166 |
| G105-D33 | G105-D33 | G105-D33 |
| G105-G31 | G105-G31 | - |
| G105-Y32 | G105-Y32 | G105-Y32 |
| G114-A44 | G114-A44 | G114-A44 |
| G149-F122 | G149-F122 | - |
| K153-T135 | K153-T135 | K153-T135 |
| - | K219-E127 | - |
| K224-S126 | - | - |
| L134-F122 | L134-F122 | L134-F122 |
| L134-S125 | L134-S125 | L134-S125 |
| L151-T135 | L151-T135 | L151-T135 |
| L151-V137 | L151-V137 | L151-V137 |
| L188-Y181 | L188-Y181 | L188-Y181 |
| L44-F101 | L44-F101 | L44-F101 |
| L44-Y88 | L44-Y88 | L44-Y88 |
| N109-H35 | N109-H35 | N109-H35 |
| N109-Q90 | N109-Q90 | - |
| N109-Y50 | N109-Y50 | N109-Y50 |
| N99-Y50 | - | - |
| P133-E127 | P133-E127 | P133-E127 |
| P133-S125 | P133-S125 | P133-S125 |
| P177-S169 | P177-S169 | P177-S169 |
| - | P177-T166 | P177-T166 |
| - | P177-T167 | - |
| Q181-E164 | Q181-E164 | Q181-E164 |
| Q61-L96 | Q61-L96 | Q61-L96 |
| R115-A44 | - | R115-A44 |
| S182-E164 | - | - |
| S187-Y181 | S187-Y181 | - |
| S189-L139 | S189-L139 | - |
| S189-V137 | S189-V137 | - |
| - | S189-Y181 | S189-Y181 |
| - | T175-Q171 | T175-Q171 |
| V179-E164 | V179-E164 | V179-E164 |
| V179-T165 | V179-T165 | - |
| V179-T166 | V179-T166 | V179-T166 |
| V191-L139 | V191-L139 | V191-L139 |
| - | - | V36-F101 |
| W113-P45 | W113-P45 | W113-P45 |

| | | |
|---|---|---|
| W46-F101<br>W46-G98<br>W46-S99<br>-<br>Y101-G51<br>Y94-A44<br>Y94-P45 | -<br>W46-G98<br>W46-S99<br>-<br>-<br>Y94-A44<br>Y94-P45 | W46-F101<br>W46-G98<br>W46-S99<br>Y101-D33<br>-<br>Y94-A44<br>Y94-P45 |

**Table S11. Average Fmax obtained from CG-SMD pulling simulations of Ab and Nb complexes with the RBD in WT, BA.4, and JN.1 variants.** Values are reported as mean ± standard deviation across 50 trajectories. For Ab/RBD WT complexes, results are shown for the complete complex (both chains) and for single-chain systems (only H or only L), highlighting the cooperative contribution of dual-chain architecture to mechanical stability.

| | Average $F_{max}$ (pN) | | |
|---|---|---|---|
| **System** | **WT** | **BA.4** | **JN.1** |
| **PDI-231 (both)** | 365.3 ± 30.1 | 472.1 ± 45.3 | 388.3 ± 42.5 |
| **PDI-231 (only H)** | 250.2 ± 25.4 | - | - |
| **PDI-231 (only L)** | 208.3 ± 31.9 | - | - |
| **S2X259 (both)** | 549.5 ± 54.2 | 472.1 ± 45.3 | 513.2 ± 61.0 |
| **S2X259 (only H)** | 427.6 ± 50.7 | - | - |
| **S2X259 (only L)** | 196.4 ± 24.8 | - | - |
| **R1-32 (both)** | 407.9 ± 45.2 | 374.9 ± 40.4 | 390.4 ± 70.4 |
| **R1-32 (only H)** | 405.2 ± 15.3 | - | - |
| **R1-32 (only L)** | 199.7 ± 20.0 | - | - |
| **R14** | 595.9 ± 55.6 | 648.6 ± 43.7 | 346.2 ± 34.4 |
| **C1** | 479.5 ± 88.2 | 513.2 ± 61.0 | 603.0 ± 55.2 |
| **n3113.1** | 430.5 ± 59.9 | 336.2 ± 74.6 | 520.7 ± 68.2 |
